# Robust Regularization Enables Automated, Real-Time Square-Wave Voltammetry Signal Quantification

**DOI:** 10.64898/2026.07.25.740173

**Authors:** Max Yates, Jenny Ji, Steven Yee, Hyongsok T. Soh

**Affiliations:** Department of Electrical Engineering, Stanford University, Stanford, CA, USA; Department of Bioengineering, Stanford University, Stanford, CA, USA; Department of Radiology, Stanford University, Stanford, CA, USA

**Keywords:** Square-wave voltammetry, electrochemical aptamer-based sensors, baseline correction, robust smoothing, regularization, real-time biosensing, signal processing

## Abstract

Square-wave voltammetry (SWV) is widely used for electrochemical biosensing because it enables sensitive, temporally resolved measurement of redox reporter signals. However, automated quantification of SWV signal remains challenging for long duration and *in vivo* measurements, where voltammograms can exhibit changing baselines, heterogeneous noise, peak drift, outliers, and interfering faradaic processes. Here, we introduce the Adaptive Square-Wave Voltammetry Iterative Fitting Toolkit (ASWIFT), an automated method for robust SWV signal extraction based on iteratively reweighted regularized smoothing. ASWIFT is available as both an open-source Python package and a downloadable desktop application. The method adaptively estimates the baseline, selects regularization strengths, fits the redox peak, and reports peak height without trace-specific parameter tuning. Across simulated datasets spanning diverse baseline, peak, noise, and concentration-response conditions, ASWIFT produced less systematic bias and more consistent signal estimates than existing methods. We further evaluated ASWIFT using *in vitro* doxorubicin measurements and *in vivo* DNA-based kanamycin sensor measurements collected in rat blood and interstitial fluid, demonstrating agreement with established methods. These results support ASWIFT as a robust framework for automated SWV analysis in real-time electrochemical biosensing.

## Introduction

Electrochemical aptamer-based (EAB) sensors offer a promising platform for the continuous monitoring of diverse molecular analytes with high sensitivity and specificity. Such sensors employ structure-switching aptamers that undergo a reversible conformational change upon binding their target. These aptamers are coupled to the surface of a gold electrode at one end and modified at the other with a redox reporter such as methylene blue. Target binding shifts the equilibrium of the aptamer away from a largely unfolded state in which the reporter is far from the electrode surface and towards a fully-folded state in which electron transfer can more readily occur between the reporter and electrode, producing a measurable current readout. EAB sensors have shown particular promise for continuous biomarker monitoring applications *in vivo* [1]. Indeed, implanted EAB sensors have enabled real-time monitoring of drugs and metabolites in live animals in experiments spanning several hours to nearly a month [2] as well as continuous drug monitoring in human subjects for 12 hours [3].

Square-wave voltammetry (SWV) is widely used to quantify the faradaic current from EAB sensors. Target binding modulates the average electron-transfer rate of the immobilized reporters, yielding a target concentration-dependent change in the SWV voltammogram. Electrochemical response is typically quantified by estimating the height of a voltammetric peak associated with the redox reporter and subtracting a baseline inferred from the background signal trace. SWV offers high sensitivity and temporal resolution, but SWV-interrogated EAB sensors also face signal-processing challenges during long-duration *in vivo* measurements in freely moving animals. *In vivo* sensors are exposed to complex biological environments that degrade the signal-to-noise ratio (SNR), induce sensor drift, and, in some cases, introduce systematic distortion from redox reactions that are unrelated to the intended reporter. These effects complicate accurate peak quantification, particularly for real-time applications requiring automated analysis. One conventional approach to address this problem is to smooth the net voltammogram using a Savitzky-Golay filter, fit the smoothed data to a high-order polynomial, and infer peak height from the extrema of the polynomial fit [4]. Although this workflow can perform well for stable datasets, it often requires empirical selection of fitting windows, smoothing parameters, and polynomial order, followed by manual inspection to identify invalid fits or refit subsets of the data. These choices may become suboptimal as the sensor drifts, the baseline changes, or transient artifacts appear, leading to peak attenuation, overfitting, or inconsistent signal measurements across an experiment.

Recent work has addressed some of these limitations through multi-Gaussian peak-baseline models enabled by automatic differentiation [5], flexible peak and baseline representations validated across diverse fitting approaches [6], and extended wavelet decomposition (EWD) to separate peak and background contributions without explicitly fitting a baseline [7]. However, these strategies still rely on user-specified parameters that may require adjustment as experimental conditions evolve. Although segmentation and periodic parameter retuning may be compatible with real-time sensing, they narrow the range of conditions under which fully automated analysis remains reliable over long durations and across varied experimental conditions. Thus, there is an unmet need for an automated analysis pipeline that generalizes across non-ideal peak shapes, baselines, and noise conditions while minimizing user-dependent analysis and parameter tuning, all while maintaining high signal precision and supporting lower limits of detection.

To address this need, we developed the Adaptive Square-Wave Voltammetry Iterative Fitting Toolkit (ASWIFT): a self-calibrating and automated approach for real-time SWV signal quantification and baseline modeling. By employing an iterative, regularization-based framework, ASWIFT provides robust smoothing and accurate peak and baseline estimates without relying on fixed parametric models, reducing the dependence on user-tuned parameters that limits previous methods. To evaluate ASWIFT, we constructed a SWV simulation pipeline that generates voltammograms with known ground-truth peak heights across diverse baseline families, additive white noise, and superimposed Gaussian peaks. In this simulation benchmark, ASWIFT produced estimates with lower systematic bias and greater consistency across replicates than prior fitting approaches, without manual optimization. We then evaluated ASWIFT on a challenging *in vitro* doxorubicin dataset, where additional faradaic processes beyond the intended redox reporter introduce substantial baseline distortions that vary across SWV frequency. Finally, we tested ASWIFT on *in vivo* kanamycin measurements from a DNA-based dual sensor, using simultaneous blood and interstitial fluid (ISF) measurements to assess performance across distinct biological sampling conditions [8]. Across these three benchmarking experiments, ASWIFT was the only method that consistently recovered signal under all conditions without changing any user-specified fitting parameters. Available as both an open-source Python package and a standalone desktop application, ASWIFT readily integrates into existing SWV analysis pipelines while also providing an accessible interface for users without programming experience. By minimizing user-dependent parameter selection, ASWIFT enables automated, accurate, and efficient real-time voltammogram analysis for SWV-based sensors, with particular utility for deriving molecular concentration estimates from *in vivo* EAB measurements.

## Results and Discussion

### ASWIFT overview

ASWIFT computes sensor response from raw voltammogram data in four steps: smoothing, baseline fitting, peak fitting, and signal calculation (**Fig. 1**). Each of the smoothing and fitting steps is formulated as a Tikhonov-regularized optimization problem [9] with adaptively selected parameters, eliminating the need for any user-dependent parameter tuning or recalibration. The key feature of ASWIFT is that the regularization strength is selected directly from each voltammogram by evaluating the tradeoff between fit fidelity and smoothness, rather than being set manually by the user. This data-driven selection is coupled with residual-based weighting, which downweights outliers and distorted regions so that parameter selection is driven primarily by the voltammetric structure rather than transient artifacts. Our method is designed for voltammograms containing a single redox peak above the baseline, while remaining tolerant of peak shouldering from secondary faradaic reactions, substantial outliers, and dynamic changes in voltammogram morphology. For voltammograms with multiple peaks, ASWIFT selects the peak with the greatest prominence by default. Within this operating regime, our method reliably identifies appropriate scaling parameters and weights across a broad range of peak shapes, noise levels, and baseline conditions. Fits are automatically rejected when no peak is detected after smoothing or when the calculated peak height is less than twice the median absolute error of the smoothed-fit residuals. These failure criteria identify cases where the redox signal is absent or difficult to distinguish from noise, improving practical usability by flagging traces that require exclusion or further inspection rather than allowing low-confidence fits to propagate through analysis unnoticed.

**Figure 1.**
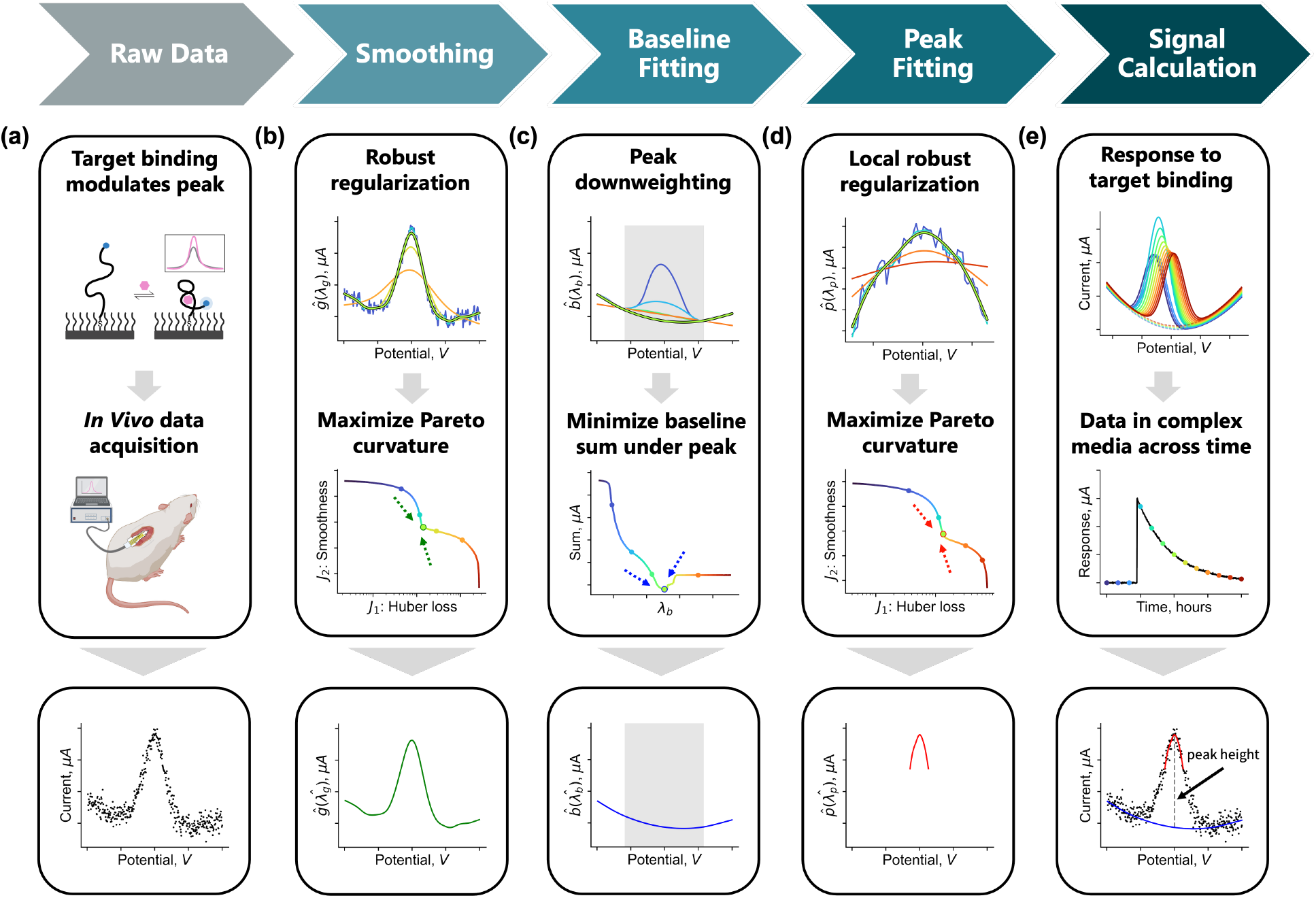
Overview of ASWIFT, an adaptive processing method for SWV data that extracts signal without requiring user-specified parameters or manual fine-tuning. **(a)** Raw SWV data is collected from an EAB sensor. Example of a raw voltammogram from an ATP-binding EAB sensor in an *in vitro* experiment. **(b)** ASWIFT first robustly denoises the raw voltammogram by balancing fidelity to the data against smoothness, while automatically downweighting outliers and selecting the regularization parameter *λ* from the maximum-curvature point of the Pareto trade-off curve. This *λ* value is then used to generate the final smoothed voltammogram. **(c)** Baseline fitting estimates a lower-envelope baseline using iterative asymmetric weighting to suppress the influence of the redox peak. The final baseline is selected as the solution that minimizes the cumulative baseline level under the automatically identified peak region. **(d)** Peak fitting applies adaptive smoothing locally within the peak region, allowing ASWIFT to evaluate candidate peak fits with different smoothness penalties and identify the locally optimal peak fit. **(e)** Changes in target concentration, sensor drift, and SWV acquisition frequency are reflected as changes in redox peak height over time. The final signal is calculated as the peak height—defined as the maximum difference between the peak fit and baseline fit—and can subsequently be normalized or further processed to infer real-time target concentration or sensor response. For **c–e**, blue curves correspond to smaller *λ* values that follow the raw data more closely, while yellow-to-red curves correspond to larger *λ* values that impose progressively stronger smoothing. The lime-green curves outlined in black at the top of panels **b–d** indicate the automatically-selected optimal solution and corresponding *λ* value.

We express the optimization problem for each step using a multi-objective cost function:

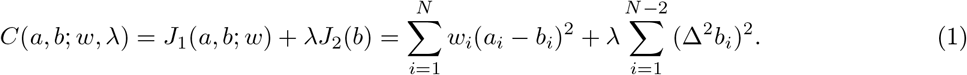

*J*_1_ is a weighted data-fidelity term that penalizes deviations between the fitted curve and the measured data, with weights *w* used to reduce the influence of undesirable points. *J*_2_ is a roughness penalty defined as the sum of squared second differences, which discourages rapid changes in curvature and enforces smoothness. The non-negative regularization parameter *λ* controls the trade-off between these objectives: smaller values emphasize fidelity to the measured data, whereas larger values emphasize smoother curvature [10]. Across steps, ASWIFT adaptively selects the *λ* value and assigns step-specific weights to balance fidelity, smoothness, and the intended fitting objective.

The smoothing step pre-treats each voltammogram before baseline fitting to reduce the risk of overfitting noise-driven deviations. This step builds on the Whittaker-Henderson smoother, which can be viewed as a form of Tikhonov regularization [11, 12]:

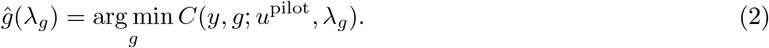

To improve robustness to heterogeneous noise, outliers, and transient artifacts, ASWIFT estimates Huber-style residual weights from a uniformly weighted pilot smooth, with residuals normalized by a 21-sample rolling median absolute deviation scale [13, 14]:

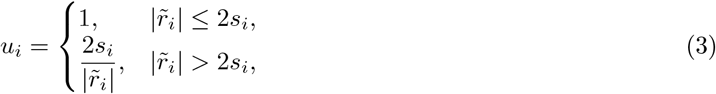

where 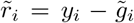 is the pilot residual, and *s*_*i*_ is the local median absolute deviation scale. These pilot weights, *u*^pilot^, downweight points that deviate strongly from the local signal structure, and are held fixed while ASWIFT solves the robust smoothing problem across a logarithmically spaced range of *λ*_*g*_ values. The final smoothing parameter, 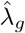, is selected using an L-curve criterion, choosing the maximum-curvature corner of the Pareto trade-off between the weighted residual objective *J*_1_ and the roughness objective *J*_2_, where additional smoothing provides diminishing noise suppression while beginning to degrade data fidelity [15, 16]. A final Huber-reweighted smoothing refinement is then performed at the selected 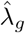.

The baseline fitting step uses the optimally smoothed signal 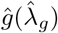 to estimate a lower-envelope baseline 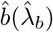.This is formulated as a second adaptively weighted smoothing problem:

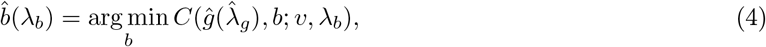

where the weights *v* are iteratively updated using the derivative peak-screened asymmetric least-squares algorithm (derpsalsa) as implemented in pybaselines [17, 18]. Conceptually, derpsalsa treats fitted points above the current baseline as likely peak contributions and exponentially downweights them. This amplitude-dependent rejection is further modulated by first- and second-derivative peak-screening terms, so that regions with peak-like slope or curvature contribute less to the baseline estimate. As a result, the fitted curve follows the lower envelope of the voltammogram while avoiding upward bias from the redox peak. For each voltammogram, ASWIFT evaluates this procedure over a logarithmically spaced range of *λ*_*b*_ values and selects the solution that minimizes the integrated fitted baseline over the peak window, where the window is defined by the peak width measured at 100% of its prominence [19]. This criterion favors a stable lower-envelope baseline that approaches linear interpolation as *λ*_*b*_ increases, while avoiding fits that overestimate the baseline beneath the redox peak.

The peak fitting step uses the same robust smoothing framework as the initial smoothing step, but restricts the fit to a local, prominence-based peak region, *P*, identified from the smoothed signal 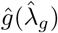. Within this region, ASWIFT estimates a smoothed peak curve 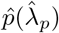 using pilot Huber weights for L-curve selection, followed by final reweighted smoothing at the selected 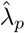:

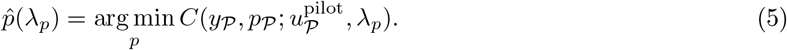

The final signal is reported as the peak height, computed as the maximum difference between the peak and baseline fits within *P*:

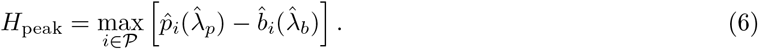

Many iterations of weighting and regularization settings can be evaluated efficiently, enabling fast, realtime SWV analysis. The resulting peak signals can then be further processed using normalization schemes [20] or kinetic differential measurements [21] to infer concentration changes or kinetics in real time with associated fitting confidence. ASWIFT also reports peak location and full-prominence peak width, with prior work demonstrating that peak location can provide useful information about sensor response [22]. Additional implementation details, including Huber-like weighting, Pareto curvature as a function of *λ*_*g*_, and derpsalsa weighting, are provided in the Methods section and in **SI, Fig. 1**.

### Simulated performance and comparison

To benchmark ASWIFT and prior methods, we implemented a simulation framework that generates voltammograms with known ground-truth peak and baseline values. Simulated data allow a direct comparison between fitted and true values that would not be possible with experimental datasets. Our simulation framework systematically varies experimental conditions by sampling across different randomly parameterized baseline families, peak widths, peak locations, and SNR values, enabling controlled and replicable evaluation of accuracy and robustness across a diverse set of simulated data. We benchmarked ASWIFT against three established SWV peak-analysis strategies: (i) polynomial peak fitting with a linearly interpolated baseline [4], (ii) multi-Gaussian symbolic fitting [5], and (iii) EWD for peak-signal isolation [7]. We did not include compiled dataset strategies such as APACE [6] in our benchmark because its performance relies on a multi-step algorithm-selection and calibration procedure that is typically tuned to a specific dataset. Incorporating this full workflow—which includes noise-level specification, peak segmentation, and algorithm screening—was outside the scope of the standardized, per-trace benchmarking framework used here. Accordingly, instead of extensively tuning each comparison method across thousands of simulated traces, we used reasonable initialization and boundary settings—including peak location, peak amplitude, and baseline parameters—that supported unsupervised fitting across the simulated dataset. In practice, more extensive tuning would require inspecting fit quality across representative traces; adjusting method-specific settings, such as voltage windows, smoothing parameters, bounds, initial guesses, or segmentation criteria; and repeating this process whenever the baseline family, peak morphology, or noise structure changes. For a benchmark containing tens of thousands of heterogeneous voltammograms, exhaustive per-condition optimization would be impractical and would effectively convert the comparison into a manually supervised workflow rather than a standardized automated benchmark. Although each method’s performance could likely be improved with additional user tuning, this sensitivity to manual parameter selection highlights a practical advantage of ASWIFT: it requires no user fine-tuning and maintains consistent performance across diverse signal and noise conditions.

We first assessed the accuracy and consistency of each method under a constant peak height of 1 *µA* using the same battery of simulated voltammograms representing different baseline families (linear, polynomial, exponential, and multi-Gaussian) and SNR levels (15–60 dB) based on peak height relative to additive white noise [23]. For each baseline family, we evaluated 30 randomly parameterized baselines for each different level of SNR. Each simulated voltammogram contained a single Gaussian peak. The peak width (defined as the 1% level relative to the amplitude) was varied from 20–50% of the sampling window. Peak location was varied between −0.25 to −0.15 V. Voltammograms were sampled at 400 points over the potential range −0.4 to 0 V, yielding traces similar to SWV EAB experimental data with a methylene blue redox reporter. Representative simulated voltammograms are shown in **Fig. 2a** for each baseline family. Further simulation details are described in Methods.

**Figure 2.**
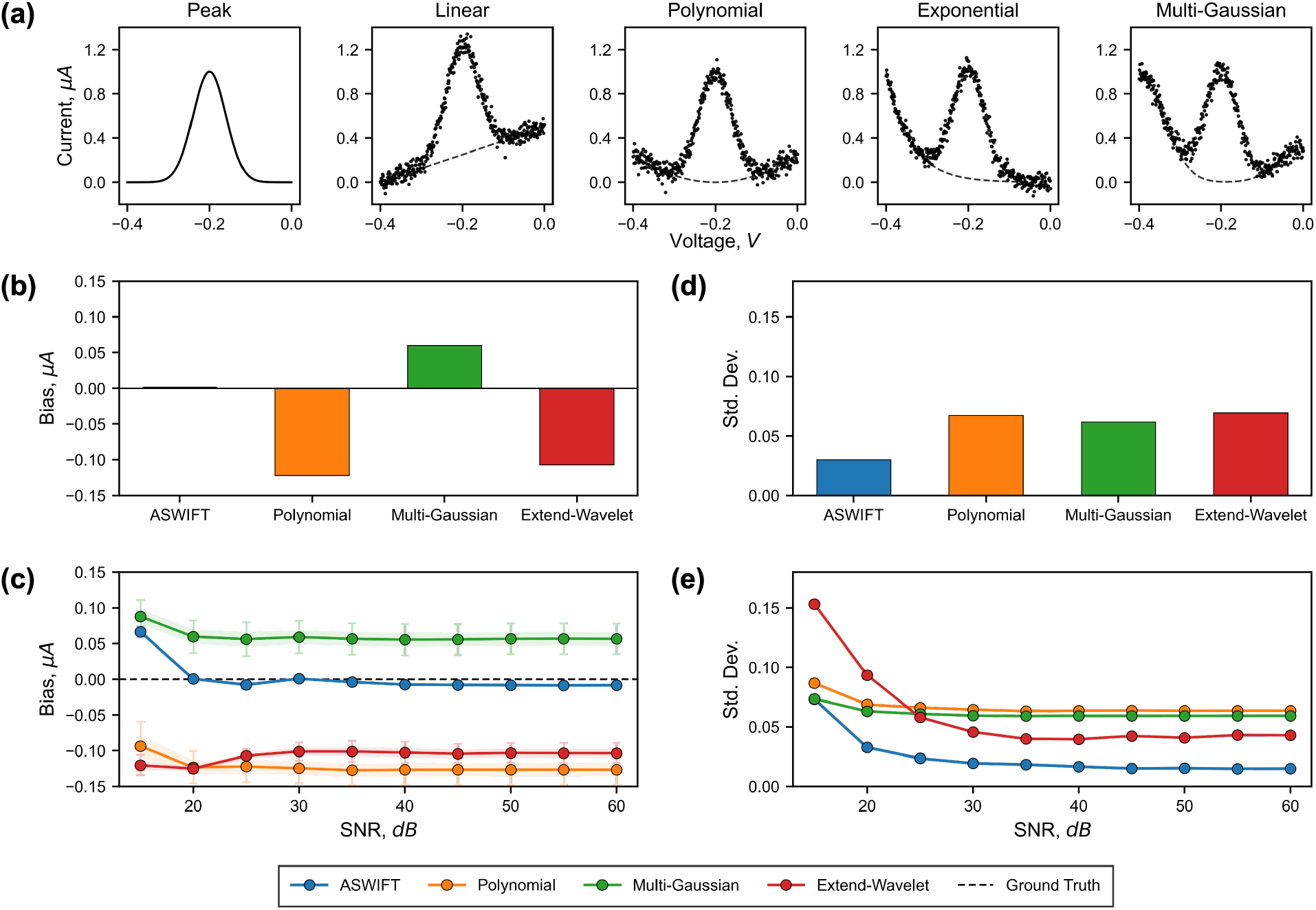
**(a)** Representative simulated voltammograms from each of the four baseline families, generated by superimposing a peak of varying width and location onto each baseline and adding Gaussian noise. **(b)** Overall bias, defined as the signed error averaged across all simulated conditions, for each fitting method. The true peak height was fixed at 1 *µA*, and values closer to zero indicate better performance. **(c)** Average bias for each method across all baseline families as a function of SNR at fixed peak height. **(d)** Overall standard deviation of the fitted peak heights across all simulated conditions. The true peak height was fixed at 1 *µA*, and lower values indicate higher precision. **(e)** Standard deviation of the fitted peak heights for each method across all baseline families as a function of SNR at fixed peak height.

To quantify accuracy, we first computed the signed error in peak height for each fit (estimated minus true). This metric quantifies method bias, defined as systematic over- or underestimation of the voltammogram peak, with an ideal value of zero. When averaged across all simulations, ASWIFT demonstrated a markedly smaller bias than other tested methods. In contrast, both the polynomial and EWD methods exhibited pronounced negative biases, indicative of systematic underestimation, whereas the multi-Gaussian method showed a positive bias, indicative of systematic overestimation (**Fig. 2b, c**). Next, we quantified consistency using the standard deviation of the fitted peak, where lower values indicate more consistent signal estimates, improved signal resolution, and a lower limit of detection [24]. Across baseline families, ASWIFT yielded the lowest standard deviation in peak height (**Fig. 2d, e**). To summarize performance across all simulated conditions, we computed the overall standard deviation across the entire dataset. By this metric, ASWIFT exhibited significantly lower variability than any other method. To further assess robustness, we repeated the simulation benchmark after introducing Laplacian-distributed outliers into a randomly selected subset of data points (**SI, Fig. 2**). This more challenging noise model produced the same qualitative performance trends observed in the primary simulations, with ASWIFT maintaining lower bias and variability than the comparison methods.

To illustrate how ASWIFT compares to other methods in estimating target concentrations, we applied each method to extract peaks from simulated datasets that mimic both signal-on and signal-off target response curves. Within each simulated dataset, the height of a Gaussian peak was set to change from 1 *µA* to a maximum or minimum of 1.5 *µA* or 0.5 *µA* for signal-on and signal-off simulations respectively. Thirteen simulated response points were generated per dataset, with peak heights associated with the expected fraction bound under Langmuir binding in the target-excess regime [24]. Target concentrations were log-spaced across four orders of magnitude centered on the two-state dissociation constant 1 *µM* . A constant level of additive Gaussian noise was applied to all voltammograms, and calibrated to achieve the specified SNR for unit-height peak. Peak location and width were held fixed, and 30 sets of baselines parameters were randomly drawn from each family.

Consistent with our first simulation, ASWIFT produced peak-height estimates closest to the ground truth for the unnormalized response, whereas the polynomial and EWD methods systematically underfit the peak and the multi-Gaussian method systematically overfit the peak (**Fig. 3a, b**). After normalizing each method to the averaged signal amplitude at 0% target fraction bound, these systematic biases were largely removed, and all four methods recovered the true binding curve on average (**Fig. 3c, d**). However, in both the unnormalized and normalized analyses, ASWIFT exhibited the lowest standard deviation in the fitted signal, thus yielding the lowest limit of detection (**Fig. 3e, f**).

**Figure 3.**
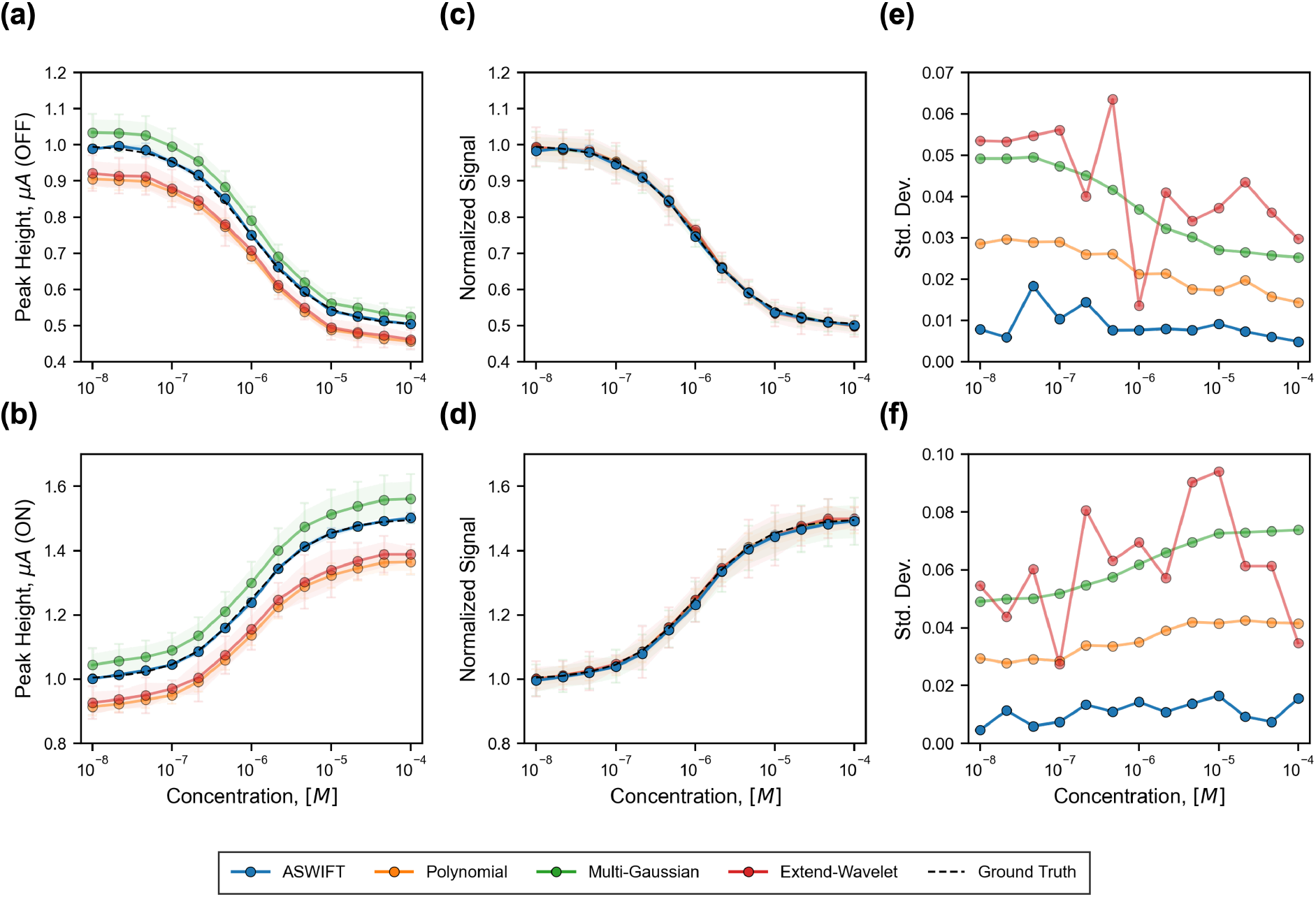
Representative simulation results showing recovery of target-response curves for each method. Points indicate the mean estimated peak height, and error bars denote the standard deviation of the fitted peak heights across randomly parameterized families of baselines. **(a, b)** Unnormalized peak-height estimates for each method applied to **(a)** signal-off and **(b)** signal-on target responses, where the true signal begins at 1 at 0% fraction bound and increases or decreases by 0.5 at 100% fraction bound. **(c, d)** Normalized peak-height estimates for each method applied to the same **(c)** signal-off and **(d)** signal-on target responses. **(e, f)** Standard deviation of the normalized signal across target concentrations for **(e)** signal-off and **(f)** signal-on sensors.

### Experimental validation

Having established ASWIFT’s accuracy and robustness in simulation, we next evaluated its performance on experimental datasets where ground truth was not directly available. We analyzed two datasets that present distinct practical challenges for SWV signal extraction. The first consisted of *in vitro* measurements collected using a doxorubicin-responsive DNA-based sensor. The second was a previously collected *in vivo* study using a DNA-based sensor to monitor kanamycin simultaneously in rat blood and ISF over several hours, with target introduced midway through the experiment [8]. Because ground truth is unavailable in these experimental settings, we did not directly quantify method-specific bias or accuracy. Instead, we assessed whether ASWIFT produced peak estimates consistent with established analytical approaches and whether it maintained stable performance under experimentally challenging baseline and noise conditions. Representative voltammogram fits for each method are provided for the *in vitro* doxorubicin datasets at low and high frequencies, the *in vivo* blood and interstitial fluid (ISF) datasets, and a challenging *in vitro* ATP dataset collected on nanoporous gold [8, 25] (**SI, Figs. 3–7**).

The *in vitro* doxorubicin dataset exhibited peak-position drift across measurements, along with substantial left-side shouldering at more negative potentials (**Fig. 4a, b**). This shoulder is likely caused by electrochemical oxidation of doxorubicin near the methylene blue redox potential [26], further challenging conventional baseline and peak-fitting methods. To evaluate concentration-dependent responses, we collected five repeat measurements across 12 independent screen-printed electrodes and two SWV frequencies, with doxorubicin concentrations ranging from 0.1 to 20 *µM* . Each trace was normalized to its corresponding electrode- and frequency-specific zero-target baseline, yielding normalized response and SNR metrics evaluated across discrete, equilibrated concentration steps (**Fig. 4c, d**). Across methods, the resulting concentration-response trends were broadly similar. At low frequency (10 Hz), however, ASWIFT was the only method that consistently maintained stable signal estimates and standard deviations at high doxorubicin concentrations (**Fig. 4e, f**), where pronounced left-side baseline shouldering relative to the methylene blue peak amplitude compromised the comparator methods. Although these other methods still returned signal estimates in this regime, the resulting values were visibly inconsistent with the true peak height and provided no internal indication that the fits were unreliable. This increases the risk of interpreting fitting artifacts as real signal changes. In contrast, ASWIFT automatically rejects unreliable fits when no valid peak can be identified or when the estimated peak is difficult to distinguish from noise, providing a practical quality-control flag rather than requiring the user to identify invalid fits through manual inspection.

**Figure 4.**
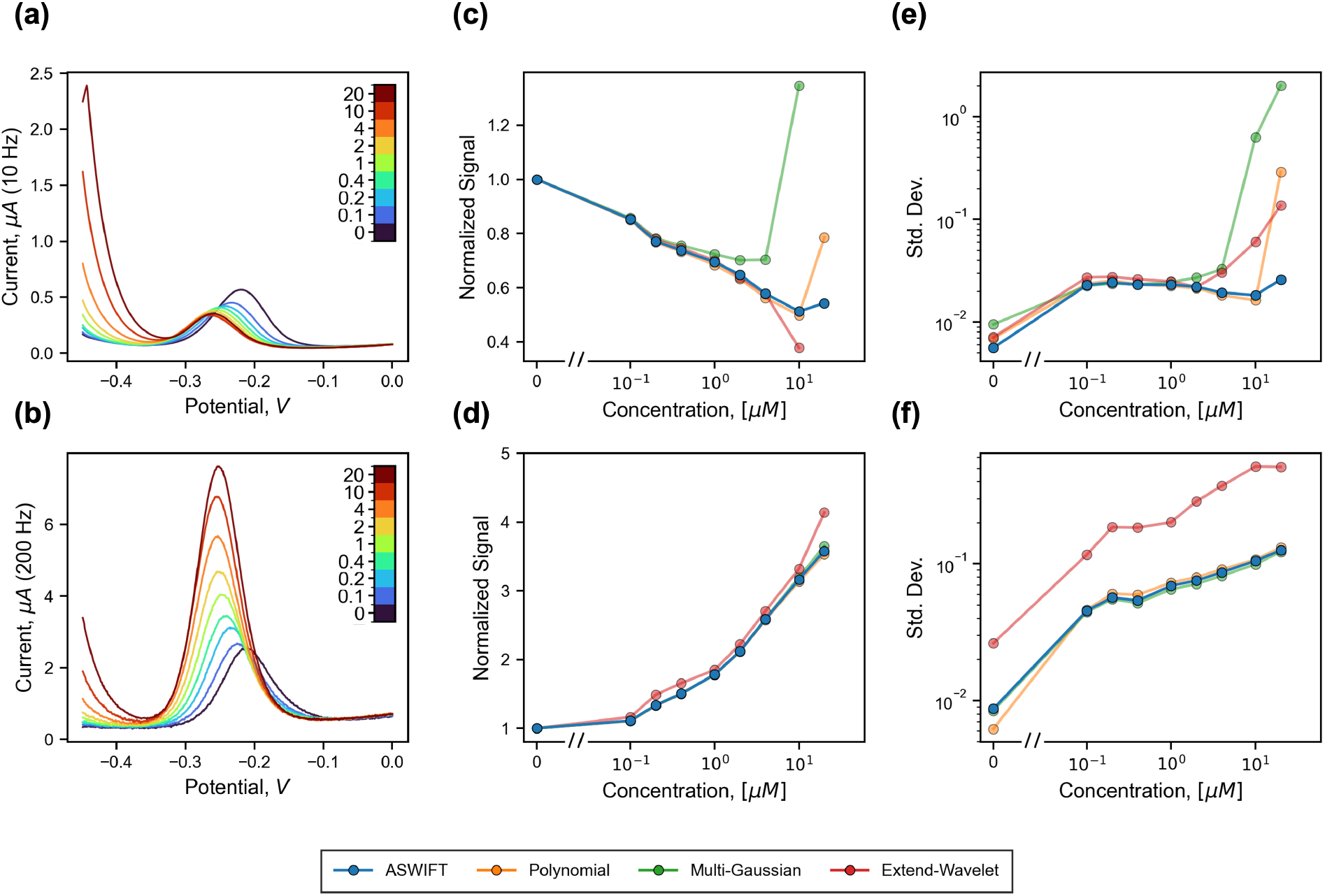
Comparing analytical methods with data from a doxorubicin-responsive DNA-based sensor. **(a, b)** Representative voltammograms collected at **(a)** 10 Hz and **(b)** 200 Hz. The “turbo” colormap encodes doxorubicin concentration in *µM* . The sensor exhibits a signal-off response at 10 Hz and a signal-on response at 200 Hz. **(c, d)** Normalized response for each method at **(c)** 10 Hz and **(d)** 200 Hz, averaged across electrodes and measurements at each concentration. High-concentration averages for comparison methods were omitted when they produced unreasonable estimates. Full plots are provided in **SI, Fig. 8. (e, f)** Standard deviation of the normalized signal across electrodes and measurements at each concentration for each method at **(e)** 10 Hz and **(f)** 200 Hz, plotted on a logarithmic scale.

The *in vivo* dataset contained voltammograms with dynamic noise, evolving baseline morphologies, and matrix-dependent differences between blood and ISF measurements, creating challenging conditions for reliable SWV signal extraction (**Fig. 5a, b**). In addition, some traces contained large transient artifacts and outliers that were accommodated by ASWIFT’s robust smoothing procedure but which impaired the other methods. Therefore, before applying the comparison methods, we used ASWIFT’s robust smoother to identify and mask points corresponding to clear artifacts, defined as measurements exceeding the smoothed signal by more than ten median absolute deviations within a local 21-sample window. Without this preprocessing step, the comparison methods failed to produce response trends consistent with ASWIFT or with the trends reported in the original study [8], which relied on manual inspection and user intervention to ensure signal fidelity. These transient artifacts likely arose from a combination of hardware instability and the challenging conditions associated with free-moving *in vivo* measurements. Additional traces and fits obtained under different outlier-filtering strengths are provided in **SI, Fig. 9**, which also demonstrates that ASWIFT’s consistency was largely independent of the filtering strength. For each method, we summarized the normalized response by computing the mean and standard deviation over consecutive, non-overlapping blocks of 10 samples (**Fig. 5c, d**). Each block spans approximately 10 minutes, over which the SWV response is approximately stationary except during target injection, although slower drift may also contribute to the measured variance. The major response trends were largely preserved across methods, excluding the multi-Gaussian method in blood due to the symbolic solver’s inability to converge to a fit close to the data given the fixed initial parameter set. However, ASWIFT maintained stronger stability in its local signal estimates in blood relative to the comparator methods (**Fig. 5e**). We attribute this difference primarily to the more challenging measurement conditions in blood, including lower SNR and more heterogeneous noise profiles. Although ASWIFT did not produce the lowest local standard deviation in ISF (**Fig. 5f**), the differences between methods were small—particularly on a logarithmic scale—indicating that multiple methods provided sufficiently stable signal estimates under these experimental conditions. Together, these results demonstrate that ASWIFT enables reliable signal extraction across distinct biological matrices without dataset-specific tuning.

**Figure 5.**
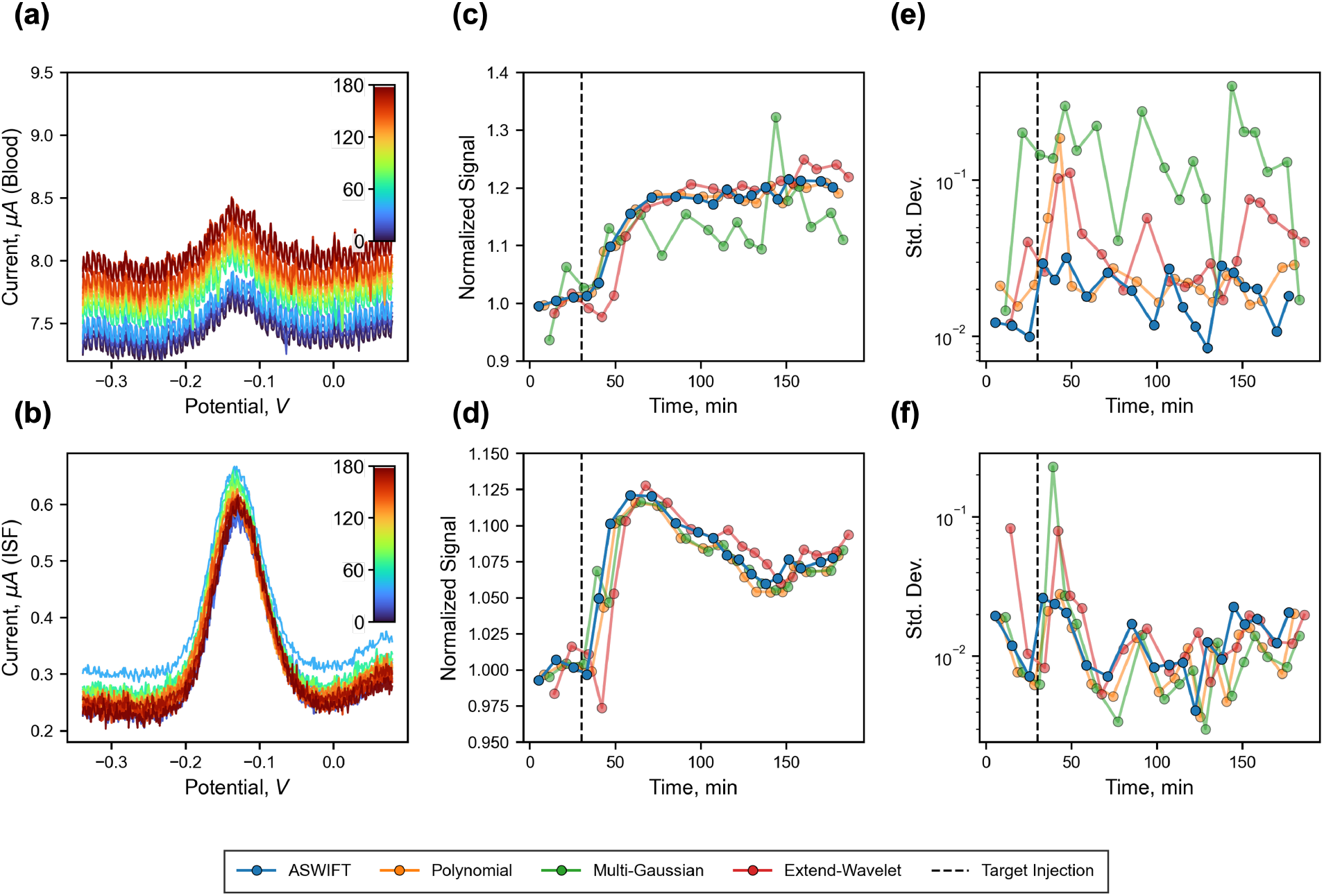
Comparing analytical methods with data from an *in vivo* rat experiment simultaneously sensing kanamycin in blood and ISF. **(a, b)** Representative voltammograms from **(a)** blood and **(b)** ISF. The “turbo” colormap encodes acquisition time on a linear scale in minutes. **(c, d)** Block-averaged, normalized response for each method for **(c)** blood and **(d)** ISF, calculated in non-overlapping windows of 10 consecutive samples. **(e, f)** Standard deviation of the normalized signal across consecutive samples for each method in **(e)** blood and **(f)** ISF, plotted on a logarithmic scale.

## Discussion

The simulation and experimental benchmarking experiments shown here demonstrate that ASWIFT provides a robust, automated framework for SWV peak quantification across diverse signal, baseline, and noise conditions. In simulations with known ground truth, ASWIFT produced lower systematic bias and lower variability than polynomial fitting, multi-Gaussian fitting, and EWD across multiple baseline families and SNR levels. This improvement was most apparent before normalization, where method-dependent bias directly affected the estimated peak height. After normalization, the average response curves from all methods became more similar, but ASWIFT still maintained the lowest signal variability. These results indicate that normalization can partially remove constant method-specific offsets, but it does not eliminate the need for robust peak extraction when comparing small signal changes, quantifying uncertainty, or operating under evolving conditions.

The experimental datasets further highlight the practical value of an analytical method that does not require trace-by-trace or dataset-specific retuning. In the *in vitro* doxorubicin dataset, peak drift and concentration-dependent shouldering created challenging fitting conditions, particularly at high target concentration and low SWV frequency. In the *in vivo* kanamycin dataset, the major response trends were largely preserved across methods, suggesting that multiple fitting approaches can recover the dominant biological signal under favorable conditions. However, the relative performance of the methods varied between blood and ISF, with ASWIFT maintaining the most consistent estimates in blood, where lower SNR and heterogeneous noise complicated signal extraction. Across these experimental conditions, ASWIFT provided the most stable overall estimates without dataset-specific retuning.

Importantly, ASWIFT should not be interpreted as a method that can recover information absent from the data or universally outperform all other approaches. With appropriate initialization, window selection, preprocessing, and dataset-specific parameter optimization, existing polynomial, multi-Gaussian, wavelet, spline, or other fitting strategies can work well for many SWV datasets. The central advantage of ASWIFT is instead operational: it reduces the manual supervision required to convert raw voltammograms into consistent peak-height estimates. Conventional fitting workflows often require users to define voltage windows, smoothing parameters, polynomial orders, initial guesses, bounds, or segmentation rules. They may also require manual inspection to identify unreliable fits and repeated retuning as peak location, peak shape, baseline curvature, SNR, or artifact structure changes. These settings may transfer across closely matched measurements from the same sensor and sample matrix, but they are not guaranteed to remain optimal across different electrodes, biological media, concentration regimes, or long-duration experiments in which drift, biofouling, or interfering redox processes alter the voltammogram over time. By contrast, ASWIFT integrates robust smoothing, adaptive regularization, lower-envelope baseline estimation, and local peak fitting into a fully automated pipeline. It also rejects low-confidence traces when no peak is detected after smoothing or when the fitted peak is indistinguishable from noise. These built-in failure criteria further reduce the need for user intervention by flagging low-confidence fits rather than allowing them to propagate through downstream analysis.

This operational advantage is closely tied to ASWIFT’s model-flexible formulation, which improves generalizability but also defines its current scope. Unlike parametric peak-fitting approaches, ASWIFT does not assume that the redox feature is Gaussian, symmetric, or described by a fixed analytical form. This flexibility is useful for EAB and related biosensors, where SWV peak shape can vary with electron-transfer kinetics, target binding, electrode condition, interfering redox-active species, and different redox reporters. However, ASWIFT is currently designed for voltammograms containing one dominant redox peak above a lower baseline. When multiple comparable peaks are present, the algorithm selects the most prominent feature by default, which may not always correspond to the chemically relevant signal. Future versions could extend the framework to explicitly handle multiple redox features, leverage baseline and peak estimates from prior measurements, or incorporate chemically informed constraints on peak identity and drift over time.

Several additional limitations should be considered. Experimental datasets do not provide ground-truth peak heights, so the experimental comparisons shown in this work primarily assess consistency and stability rather than absolute accuracy. In addition, the benchmarking was performed using reasonable implementations of existing methods rather than exhaustive user-specific optimization of every parameter. This choice reflects the intended real-time use-case across many different experiments, but it also means that the comparisons should be interpreted as practical pipeline-level benchmarks rather than definitive limits on each method’s best possible performance. Finally, accurate concentration inference from SWV peak heights still depends on sensor calibration, normalization strategy, binding kinetics, and drift correction beyond the signal extraction step itself.

Overall, ASWIFT addresses a key bottleneck in SWV-based biosensing: reliable, automated conversion of raw voltammograms into quantitative peak-height estimates under nonideal conditions. While conventional signal-processing methods can often be tuned to perform well for a specific sensor and sensing environment, this tuning may need to be revisited as electrode fabrication, sensor loading, matrix composition, fouling, baseline drift, SNR, and artifact structure vary across experiments or over the course of continuous measurements. By reducing dependence on user-defined fitting parameters while maintaining strong performance across simulations and experimental datasets, ASWIFT enables more consistent analysis of electrochemical measurements without repeated dataset-specific optimization. This capability is particularly important for continuous EAB sensing, where real-time concentration tracking requires automated processing of large numbers of voltammograms acquired in complex biological environments.

## Methods

### Regularizer implementation

We write the weighted Tikhonov optimization problem in Eq. 1 in matrix form as

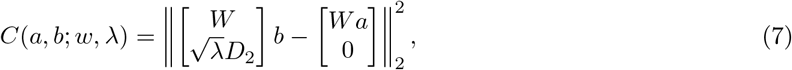

where *a, b* ∈ *R*^*n*^ denote the measured and fitted signals, respectively, and *w* ∈ *R*^*n*^contains the residual weights. Here, 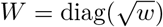 is the diagonal weighting matrix and *D*_2_ ∈ *R*^(*n*−2)×*n*^ is the second-difference matrix, defined such that

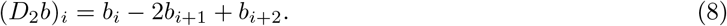

For fixed weights and regularization parameter *λ*, the minimizer is

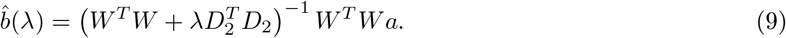

Because *W*^*T*^ *W* is diagonal and 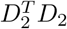 is pentadiagonal, this system can be solved efficiently as a sparse pentadiagonal linear system with linear computational complexity in the number of sampled data points [19].

For the initial smoothing and peak-fitting steps, ASWIFT first computes a uniformly weighted pilot smooth, with the pilot regularization parameter selected from the maximum-curvature corner of the unweighted L-curve. The resulting residuals are then used to estimate Huber-style weights. Residuals are normalized by a local scale computed as a 21-sample rolling median absolute deviation, and points with residuals larger than two local scale are progressively downweighted according to Eq. 3. These pilot weights are held fixed during the weighted L-curve sweep used to select *λ*_*g*_ or *λ*_*p*_. After the optimal regularization parameter is selected, a final Huber-reweighted smoothing refinement is performed at the selected *λ*. For the peak-fitting step, smoothing was restricted to the local peak window, defined as the prominence-based width of the largest identified peak at a relative height of 0.3 [19]. Optimization of the static parameters held fixed across all datasets, including the Huber local-scale cutoff, rolling window size, and relative-height threshold for peak-region identification, is shown in **SI, Figs. 10-12**.

The regularization parameter is selected from a logarithmically spaced grid using an L-curve criterion. For each candidate *λ*, ASWIFT evaluates the weighted residual objective *J*_1_ and the roughness objective *J*_2_, then selects the point of maximum signed planar curvature along the log-objective Pareto curve,

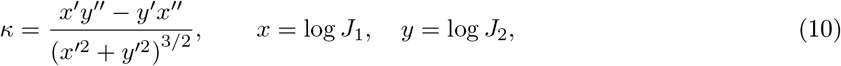

where primes denote finite differences along the logarithmically sampled *λ* grid [15, 16].

### Simulation details

Simulated voltammograms were generated as the sum of a known baseline, a single Gaussian peak, and additive Gaussian noise:

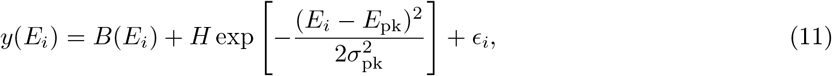

where *E*_*i*_ is the sampled potential, *B*(*E*_*i*_) is the true baseline, *H* is the true peak height, *E*_pk_ is the peak location, and 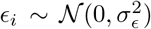 is independent additive noise. The four families of baselines were generated using a fixed pseudorandom seed and were constrained so that they did not introduce a second redox-like peak. The baseline models and parameter ranges are summarized in **Table 1**. The simulation grids used for the constant-amplitude benchmark in **Fig. 2** and target-response benchmark in **Fig. 3** are summarized in **Table 2**.

**Table 1:**
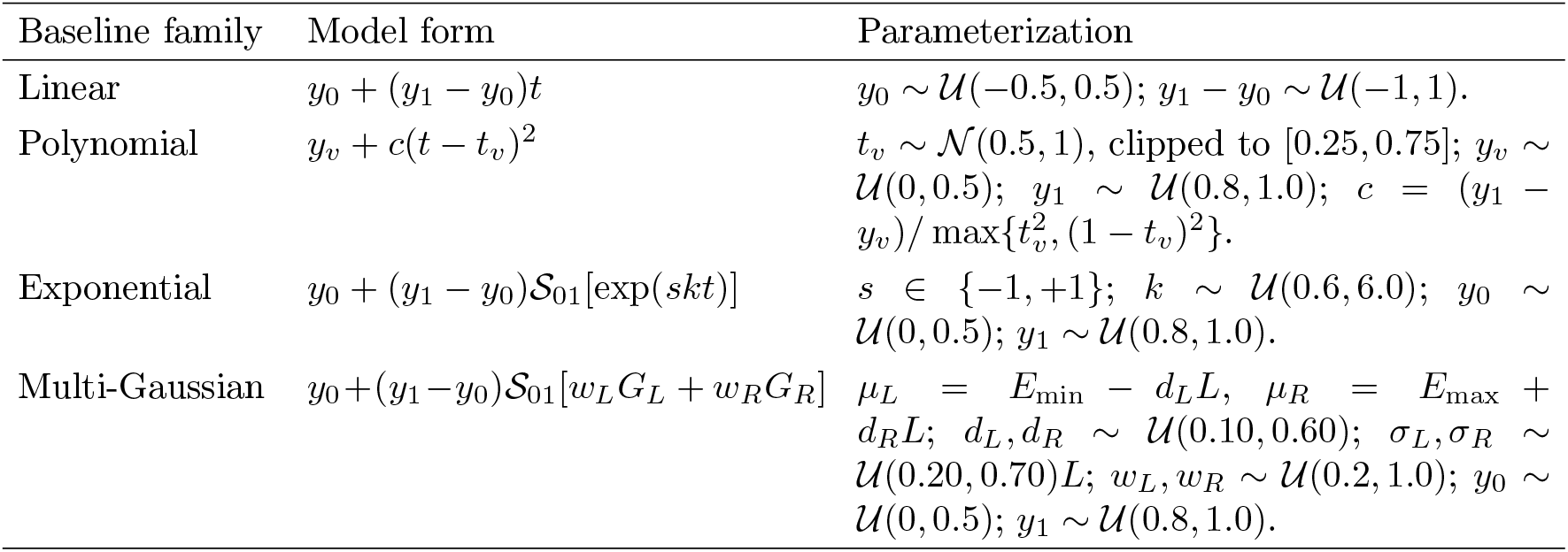
Baseline families used for simulated voltammograms. Here, *t* = (*E* − *E*_min_)/(*E*_max_ − *E*_min_) is the normalized voltage coordinate, with *t* = 0 at *E*_min_ and *t* = 1 at *E*_max_, and *L* = *E*_max_ − *E*_min_ is the voltage-window width. For any sampled function *q*(*E*), *S*_01_[*q*] = (*q*− min *q*)/(max *q* − min *q*) denotes minmax normalization over the sampled voltage window, rescaling *q* to the range [0, 1]. *G*_*L*_ = *G*(*E*; *µ*_*L*_, *σ*_*L*_) and *G*_*R*_ = *G*(*E*; *µ*_*R*_, *σ*_*R*_) denote the left and right Gaussian shoulder components.

**Table 2:** Simulation grids used for **Fig. 2** and **Fig. 3**.

| Parameter | Fig. 2 benchmark | Fig. 3 benchmark |
| --- | --- | --- |
| Potential range | −0.4 to 0 V | −0.4 to 0 V |
| Sampling | 400 evenly spaced points | 400 evenly spaced points |
| Baseline panel | 4 families; 30 shapes per family | 4 families; 30 shapes per family |
| Peak model | Single Gaussian | Single Gaussian |
| Peak height | Fixed at 1 $\mu\text{A}$ | On: $1 + 0.5f_b$; Off: $1 - 0.5f_b$ |
| Peak location | −0.25, −0.20, −0.15 V | −0.20 V |
| Peak width | 20%, 30%, 50% of window | 30% of window |
| SNR | 15–60 dB; 5 dB spacing | 30 dB |
| Noise replicates | 1 | 3 |
| Total traces | 10,800 | 4,680 signal-on; 4,680 signal-off |

Peak width was defined by the distance over which the Gaussian decayed to 1% of its peak amplitude. For a fractional width *w* of the voltage window, *σ*_pk_ was selected such that

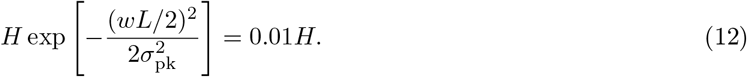

Noise levels were set by the peak signal-to-noise ratio,

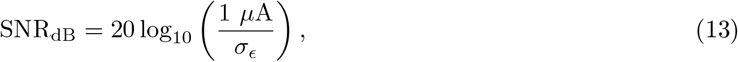

where the signal amplitude was defined relative to the 1 *µ*A reference peak height. For the **Fig. 3** target-response simulations, the noise amplitude was fixed at the value corresponding to a 30 dB SNR relative to a 1 *µA* peak height.

For the constant-amplitude simulations in **Fig. 2**, all combinations of baseline family, baseline parameterization, SNR, peak width, and peak location were evaluated, yielding 4 × 30 × 10 × 3 × 3 = 10,800 simulated voltammograms. For the target-response simulations in **Fig. 3**, the true peak height was varied to represent idealized signal-on and signal-off responses:

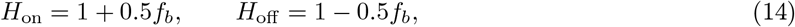

where *f*_*b*_ is the simulated fraction bound. Thirteen fraction-bound values were used, spanning approximately 1% to 99% bound:

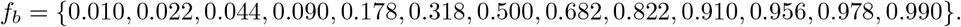

Three independent noise replicates were generated for each amplitude and baseline class, yielding 4 × 30 × 13 × 3 = 4,680 traces for each of the signal-on and signal-off simulations. Normalized response curves were calculated by dividing the fitted peak heights by the mean fitted peak height at 0% fraction-bound for each method.

### Doxorubicin *in vitro* experiment

DNA with sequence /5ThioMC6-D/AC CAT CTG TGT AAG GGG TAA GGG GTG GT/3AmMo/ bearing a 5’ thiol modifier C6 S-S and a 3’C6 amino modifiers was synthesized in-house (Biolytic Expedite 8909, Glen Research phosphoramidites), labeled using ATTO-MB2 NHS ester (Leica), and purified by ethanol precipitation followed by reverse phase HPLC.

DNA solutions for electrode immobilization were prepared by reduction with a 1,000-fold molar excess of Tris(2-carboxyethyl)phosphine (TCEP) in water (66 *µM* DNA and 66 *mM* TCEP) for 90 min. The reduced DNA was then diluted to 500 *nM* in a final buffer condition containing 1x phosphate-buffered saline (from 10x PBS, Fisher BP3991) with an additional 2 *M* NaCl.

Twelve independent screen-printed electrodes (Metrohm 220BT) were electrochemically cleaned using cyclic voltammetry (− 0.4 to 1.4 *V* at 300 *mV/s*) in 0.5 *M* sulfuric acid until the reduction peak measured − 0.6 *mA* using a multi-channel potentiostat (MultiEmstat4, PalmSens). The electrodes were then rinsed with MilliQ water, dried under nitrogen stream, and incubated with 40 *µL* of the reduced 500 *nM* DNA solution for 5 h. Electrodes were then rinsed with 1x PBS, gently and briefly dried under nitrogen stream, and incubated overnight in a 100 *µL* 5 *mM* 6-mercapto-1-hexanol solution in 1x PBS. Finally, the electrodes were gently rinsed with 2x selection buffer (10 *mM* Tris-HCl pH 8, 60 *mM* NaCl, 2.5 *mM* KCl, 0.5 *mM* MgCl_2_, 0.5 *mM* CaCl_2_, 0.005% v/v Tween-20) and briefly dried under nitrogen stream, and equilibrated in 2x selection buffer for 2 h before testing.

Prepared electrodes were subsequently subjected to SWV interrogation (-0.45 to 0 *V* at 10, 100, 200, 500, 700 Hz) in a range of doxorubicin concentrations (0, 0.1, 0.2, 0.4, 1, 2, 4, 10, 20 *µM*) prepared in 2x selection buffer. For each concentration, a set of five replicate measurements was taken for each electrode.

## Supporting information

Extended simulations and data analysis, trace-by-trace fit comparisons, optimization of static parameters.

Instructions for installing, configuring, and using ASWIFT.

## Abbreviations

ASWIFT: Adaptive Square-Wave Voltammetry Iterative Fitting Toolkit
SWV: square-wave voltammetry
EAB: electrochemical aptamer-based
SNR: signal-to-noise ratio
ISF: interstitial fluid
EWD: extended wavelet decomposition
APACE: algorithm-powered analyzer for continuous electrochemistry
KDM: kinetic differential measurement
MB: methylene blue
ATP: adenosine triphosphate
LOD: limit of detection.

## Acknowledgments

The authors thank Kevin Plaxco and Zeki Duman for sharing *in vivo* ISF and plasma datasets for doxorubicin, tobramycin, and cocaine sensing [27, 28], which were used to further validate ASWIFT but are not included in this manuscript. The authors thank Ya-Chen Tsai for helpful discussion on implementing extended wavelet decomposition [7]. The authors thank Yihang Chen for sharing the *in vivo* data we used in **Fig. 5** [8]. The authors thank Michael Eisenstein for editing this manuscript and Linus Hein for helpful discussions on multi-objective optimization and efficient numerical implementation.

## Author contributions

- Conceptualization: M.Y., J.J., S.Y.
- Methodology: M.Y.
- Investigation: M.Y., J.J., S.Y.
- Visualization: M.Y., J.J., S.Y.
- Funding acquisition: H.T.S.
- Project administration: H.T.S.
- Writing–original draft: M.Y., H.T.S.
- Writing–reviewing & editing: all authors

## Data availability

The ASWIFT source code is available at: https://github.com/Soh-Lab/aswift. This repository also contains a ZIP archive of all raw experimental data in the release assets. The simulated datasets and code used to generate all figures are available at: https://github.com/Soh-Lab/aswift_figures.

## Supporting information

The following files are available free of charge:

- ASWIFT_SI.pdf: Extended simulations and data analysis, trace-by-trace fit comparisons, optimization of static parameters.
- ASWIFT_UG.pdf: Instructions for installing, configuring, and using ASWIFT.

