## Extended simulations and data analysis, trace-by-trace fit comparisons, optimization of static parameters. for "Robust Regularization Enables Automated, Real-Time Square-Wave Voltammetry Signal Quantification"

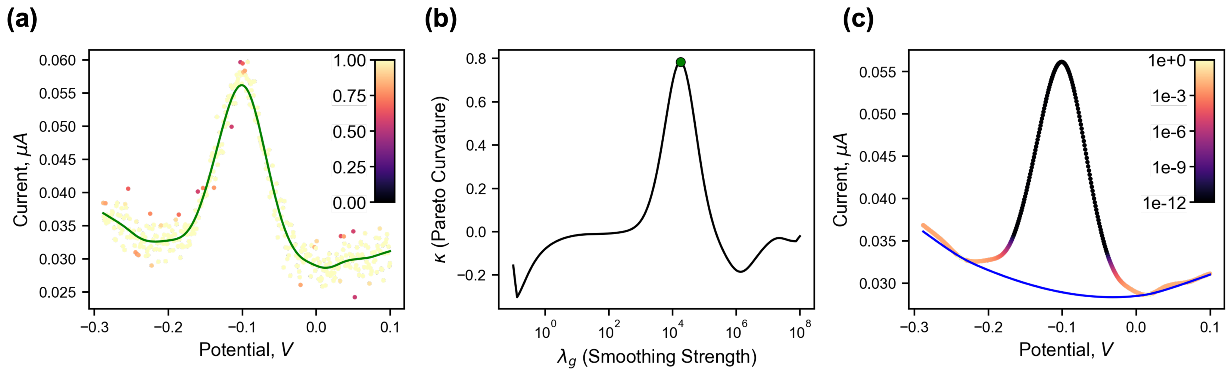


**Figure S1:** Weighting behavior and regularization-parameter selection for the smoothing and baseline-fitting steps shown in **Fig. 1**. **(a)** Final Huber-like weights used during robust smoothing, overlaid on the raw voltammogram and smoothed signal. Weights are shown on a linear scale in the inset colorbar. **(b)** Pareto curvature, $\kappa$, computed from the tradeoff between the data-fidelity $(J_1)$ and roughness penalty $(J_2)$ during smoothing, as defined by Eq. 10. The selected regularization parameter corresponds to the maximum signed-curvature point. **(c)** Final derpsalsa weights used during adaptive baseline fitting, overlaid on the smoothed voltammogram and fitted baseline. Weights are shown on a logarithmic scale in the inset colorbar.

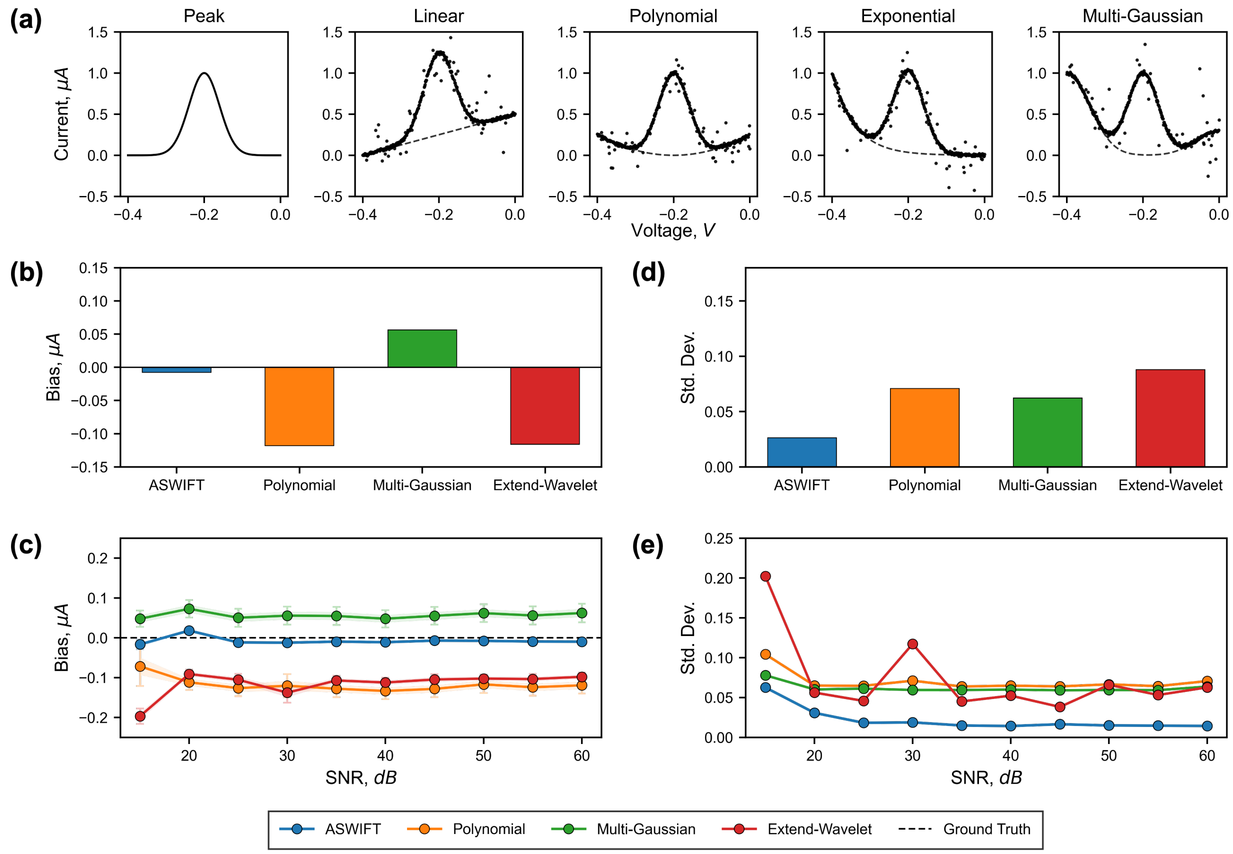


**Figure S2:** Performance of each method after random injection of outliers. **(a)** Representative simulated voltammograms from each of the four baseline families. We used the same simulation panel from **Fig. 2**, with additional Laplacian noise injected into a uniformly sampled random subset comprising 0–20% of the data points in each simulated trace. The Laplace scale parameter was fixed at $b = 0.12$ for all traces, independent of SNR. **(b)** Overall bias, with the true peak height fixed at 1 $\mu A$. **(c)** Average bias for each method as a function of SNR. **(d)** Overall standard deviation of the fitted peak heights across all simulated conditions. **(e)** Standard deviation of the fitted peak heights as a function of SNR.

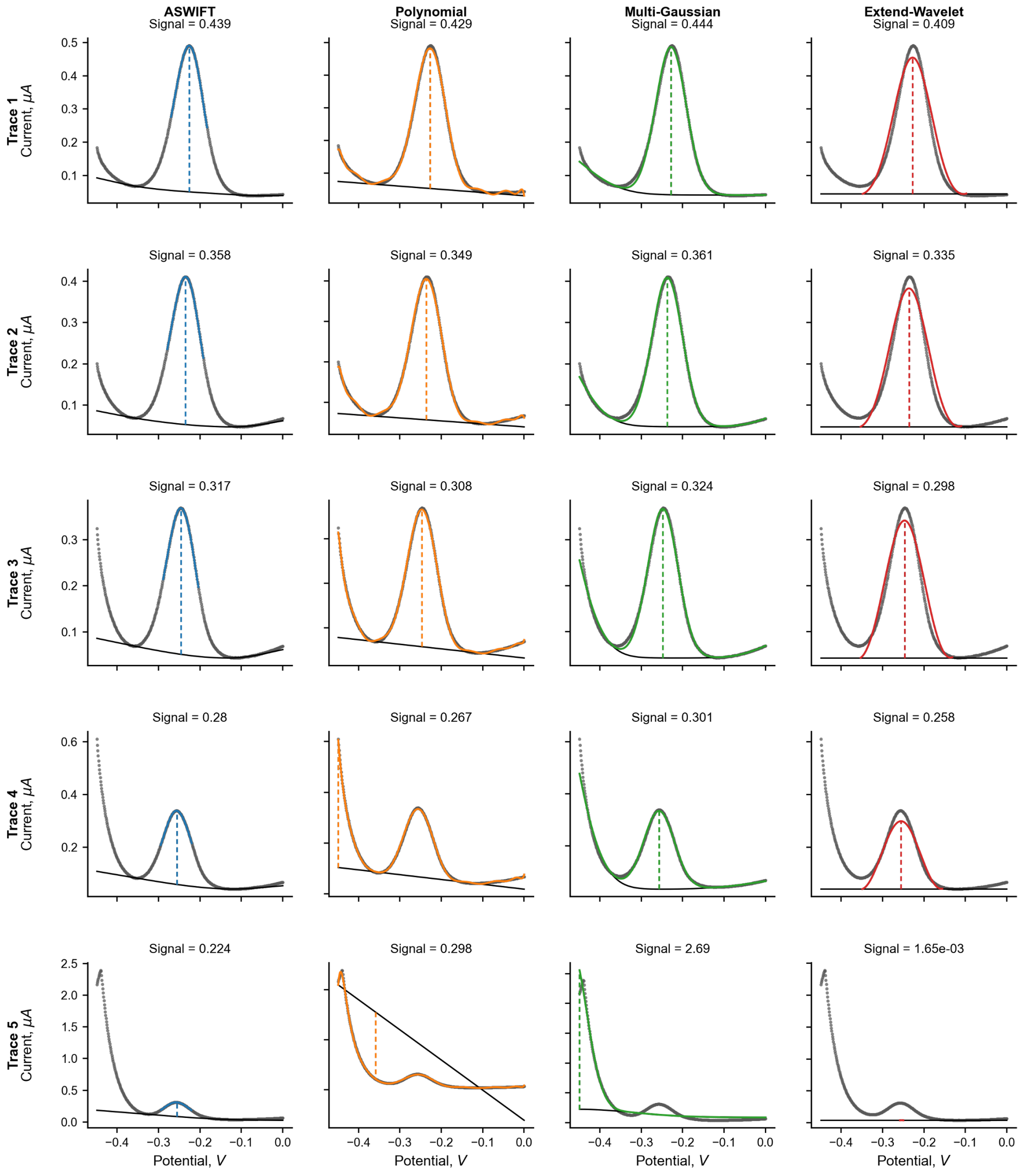


**Figure S3:** Representative fits to selected *in vitro* doxorubicin voltammograms measured in 2x selection buffer at 10 Hz from the dataset analyzed in **Fig. 4**. Five representative traces are shown for each method. For methods that explicitly fit a baseline and peak model, the plotted curves correspond to the fitted baseline, fitted peak, and reconstructed signal. For EWD, which does not explicitly estimate a baseline, we visualized the result by plotting the denoised wavelet reconstruction as the peak contribution and a constant baseline defined by the minimum value of the smoothed curve on either side of the peak region identified at a relative-height prominence of 1.0. Across methods, peak localization was more challenging in the presence of pronounced left-side voltammogram shouldering.

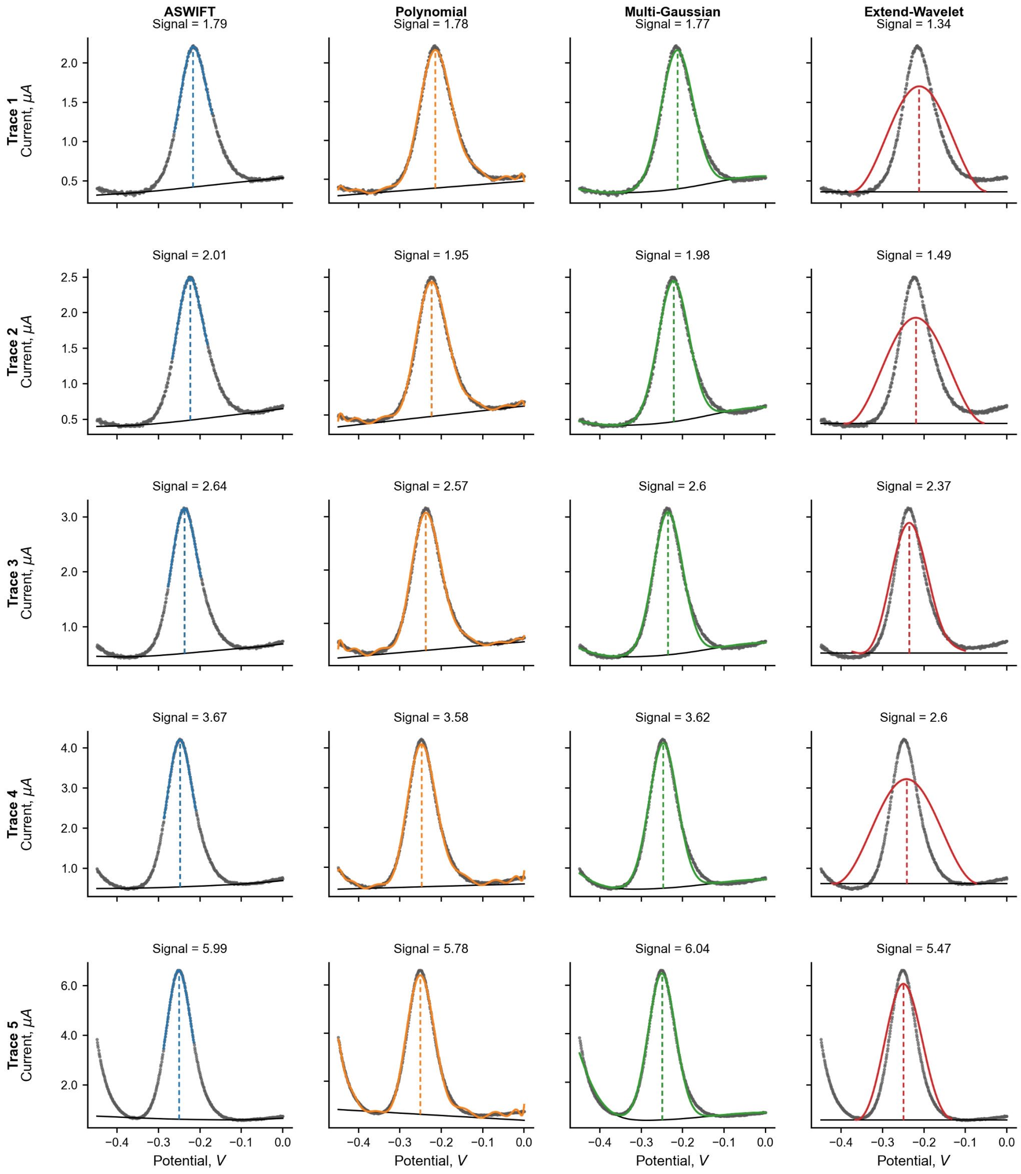


**Figure S4:** Representative fits to selected *in vitro* doxorubicin voltammograms measured in 2x selection buffer at 200 Hz from the dataset analyzed in **Fig. 4**. Five representative traces are shown for each method using the plotting conventions described in **SI, Fig. 3**. In these examples, EWD tended to produce broader, lower peak reconstructions, most likely because baseline contributions with peak-like morphology were incorporated into the denoised reconstruction.

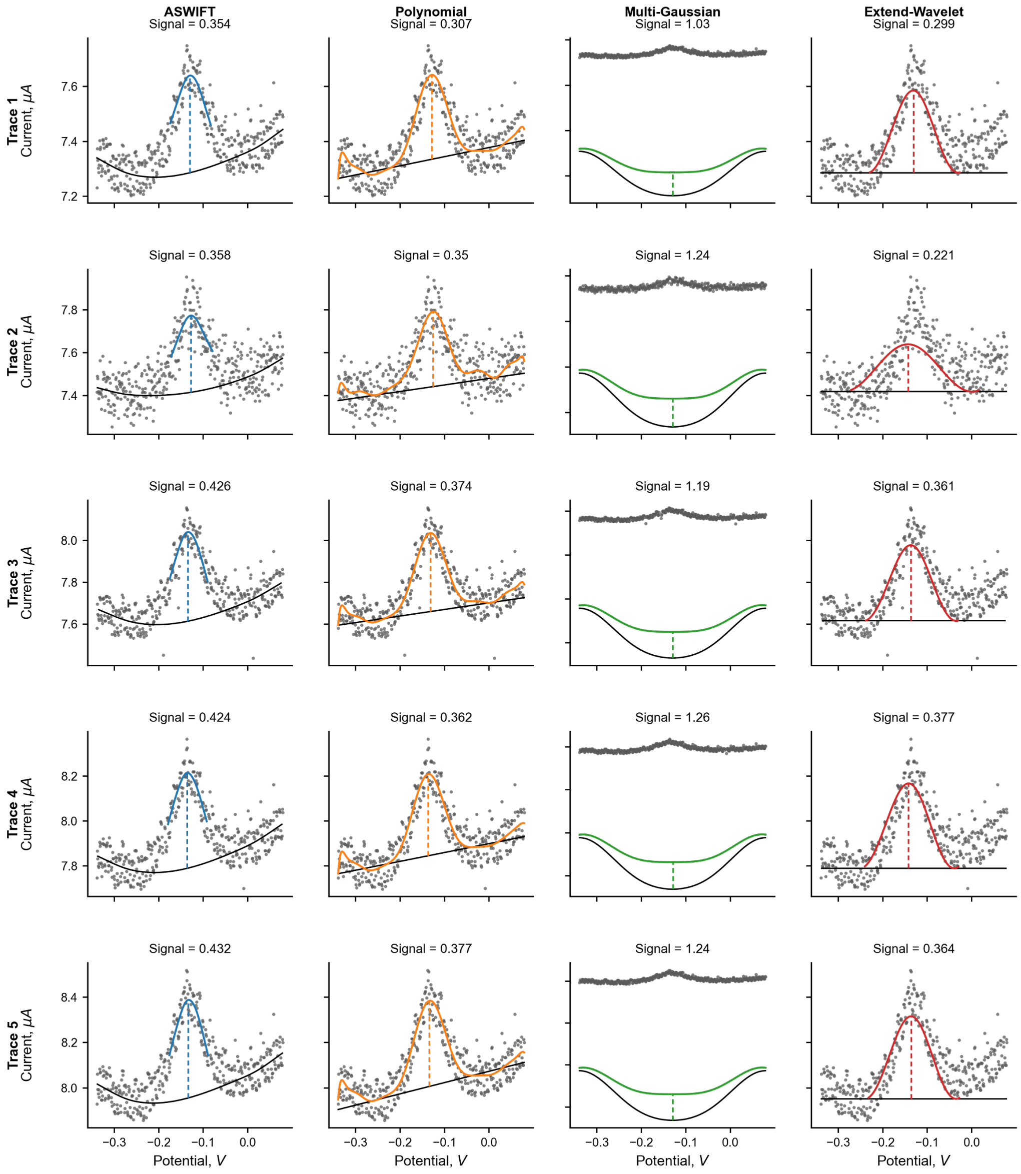


**Figure S5:** Representative fits to selected *in vivo* kanamycin voltammograms measured in blood from the dataset analyzed in **Fig. 5**. Five representative traces are shown for each method using the plotting conventions described in **SI, Fig. 3**. Across these examples, Multi-Gaussian fits converged to parameter estimates that did not closely match the measured voltammogram, while EWD tended to produce broader, lower peak reconstructions when noise was concentrated near the baseline or voltammogram tails.

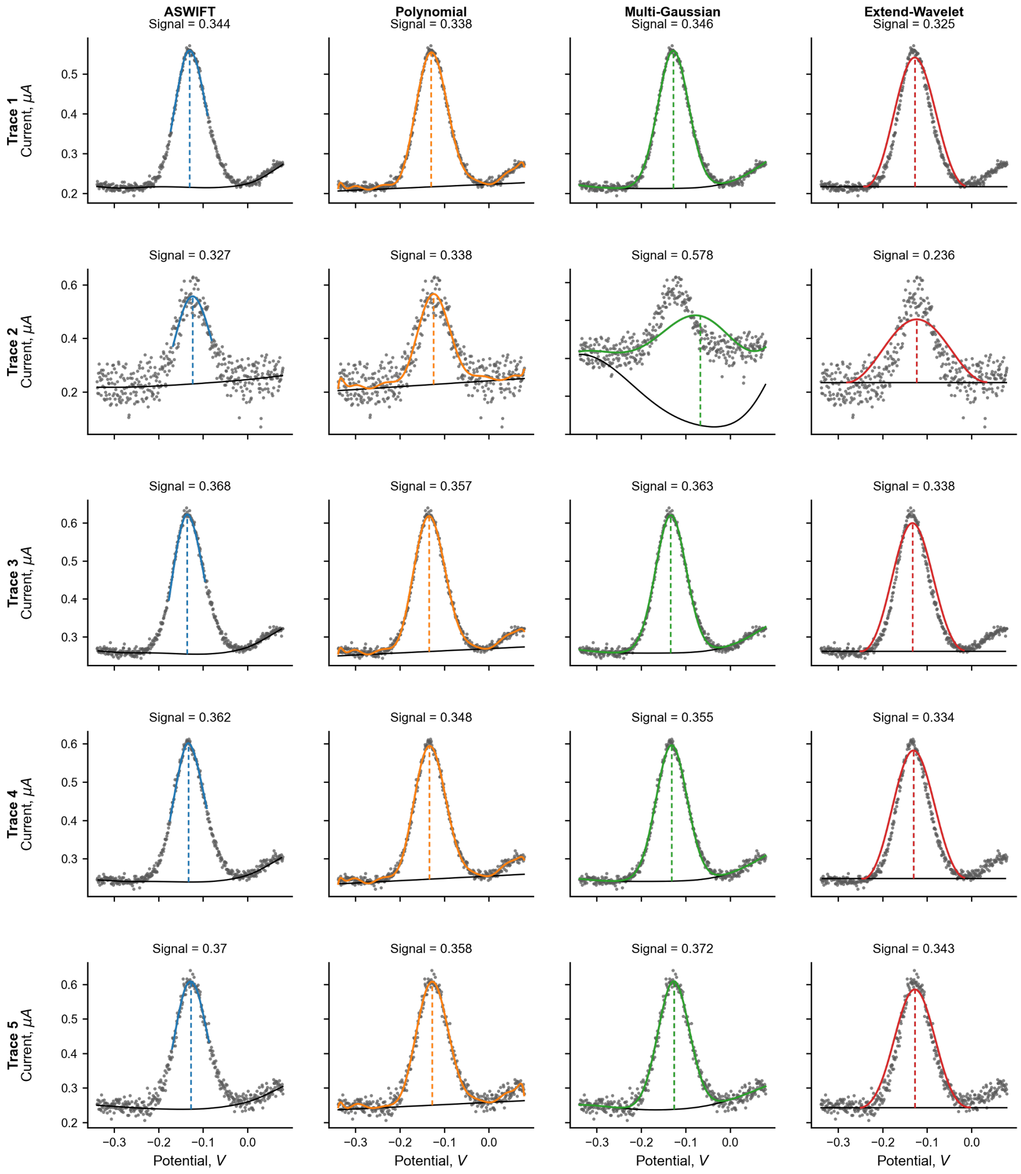


**Figure S6:** Representative fits to selected *in vivo* kanamycin voltammograms measured in ISF from the dataset analyzed in **Fig. 5**. Five representative traces are shown for each method using the plotting conventions described in **SI, Fig. 3**. These examples show similar fitting behavior to the blood dataset, including occasional multi-Gaussian fits that did not closely match the measured voltammogram, and broader EWD peak reconstructions when noise was concentrated near the baseline or voltammogram tails.

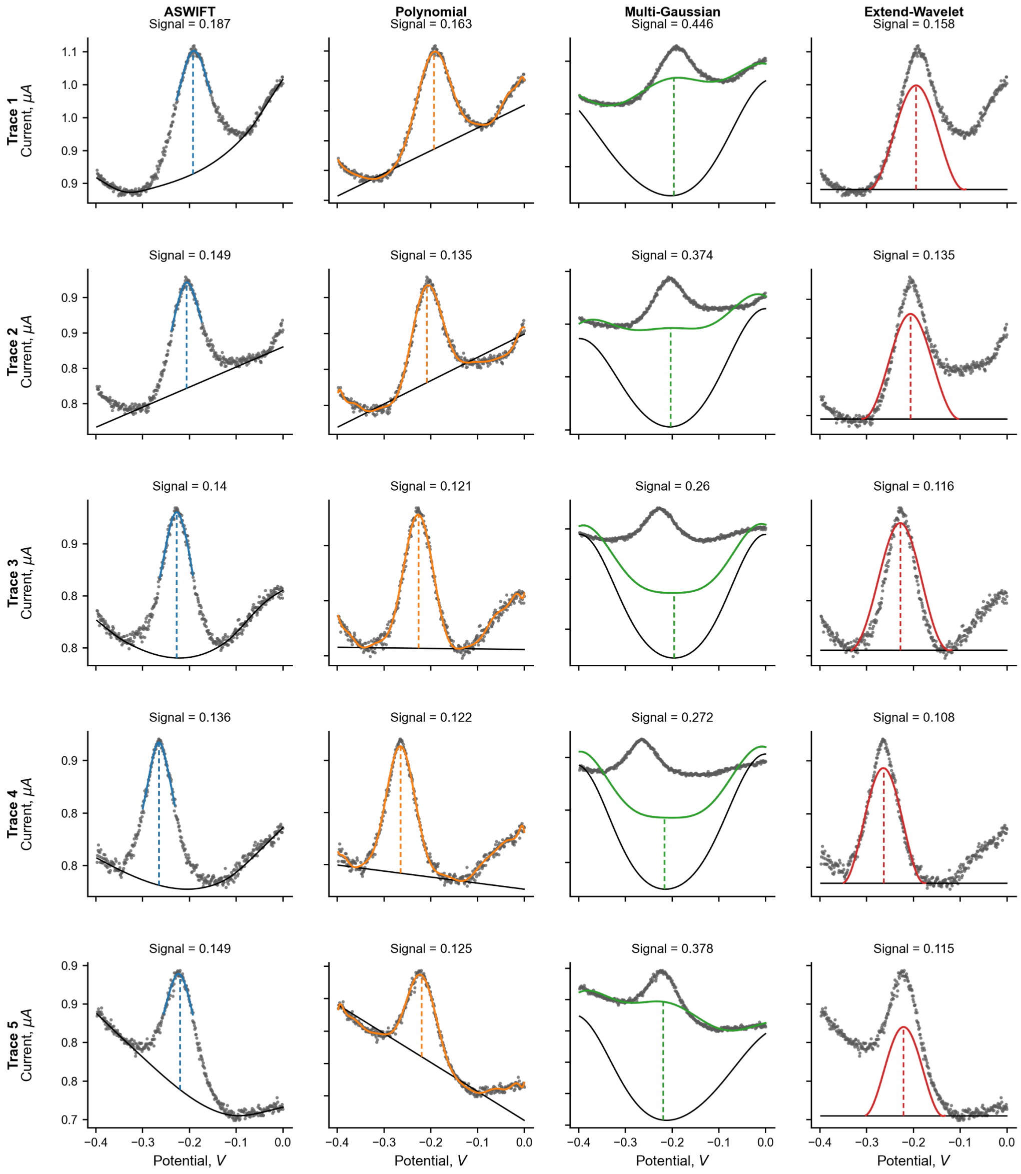


**Figure S7:** Representative fits to selected *in vitro* ATP voltammograms measured from an EAB aptamer sensor immobilized on a suboptimal nanoporous gold surface and tested in 2x selection buffer at 150 Hz. Five representative traces selected from a binding curve series are shown for each method using the plotting conventions described in **SI, Fig. 3**. In these examples, several method-dependent fitting behaviors were observed, including polynomial fits that interpolated the baseline through an inaccurate point without precise user intervention, multi-Gaussian fits that did not closely match the measured voltammogram, and broader EWD peak reconstructions.

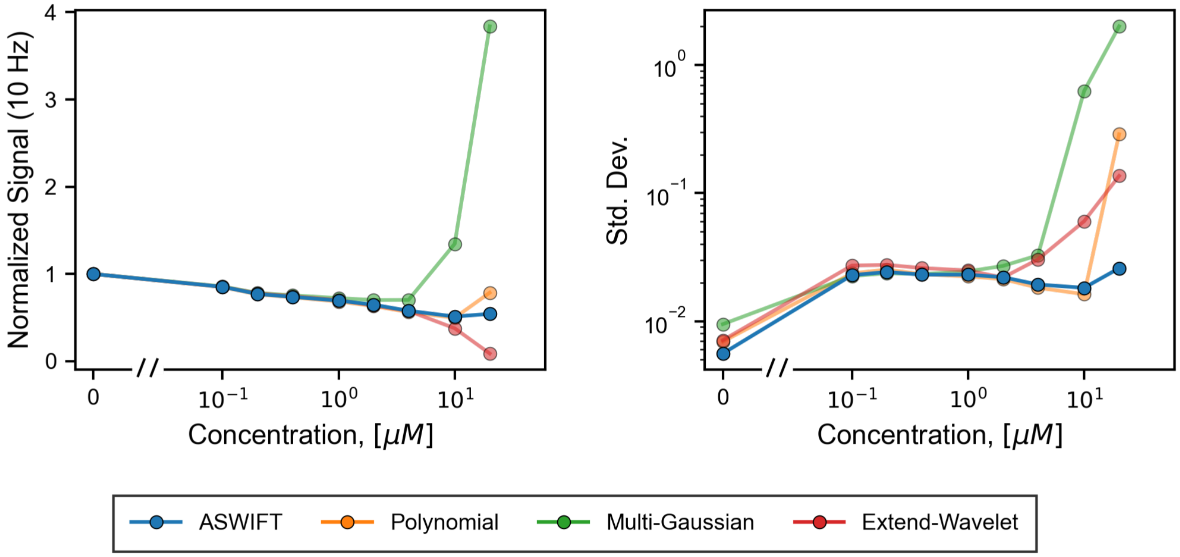


**Figure S8:** Full plots corresponding to the first row of **Fig. 4**. These panels show the complete high-concentration responses, including regions that were partially omitted for other methods in **Fig. 4** because inaccurate peak estimation distorted the reported signal.

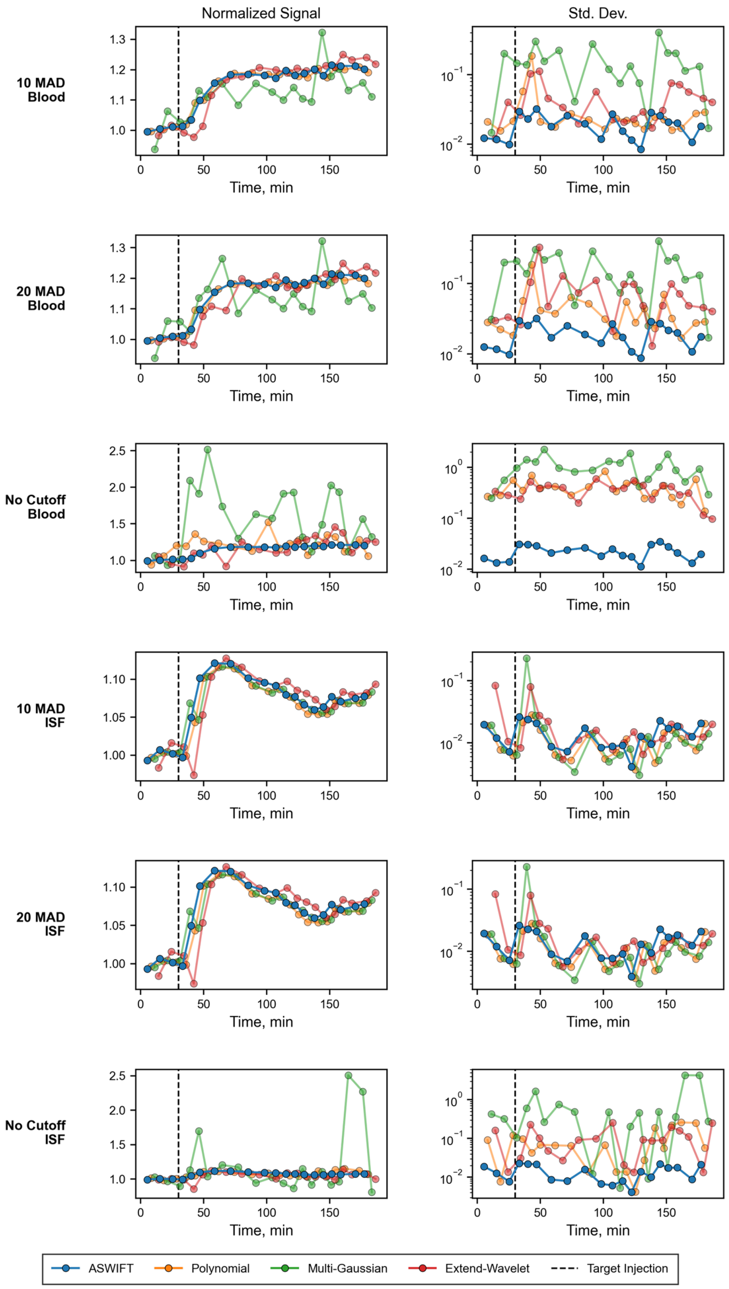


**Figure S9:** Normalized response and standard deviation calculated over 10 consecutive samples for the *in vivo* blood and ISF experiments. Three median absolute deviation cutoff thresholds are shown. A 10x cutoff, which was used in **Fig. 5**, reduced the influence of large transient artifacts while preserving comparable response trends across methods. Increasing the cutoff to 20x increased the standard deviation for the other methods, particularly in blood, whereas omitting the cutoff substantially increased the standard deviation in both blood and ISF. Because these datasets were affected by faulty hardware and challenging *in vivo* measurement conditions, we applied the 10x cutoff to enable more consistent comparison across methods.

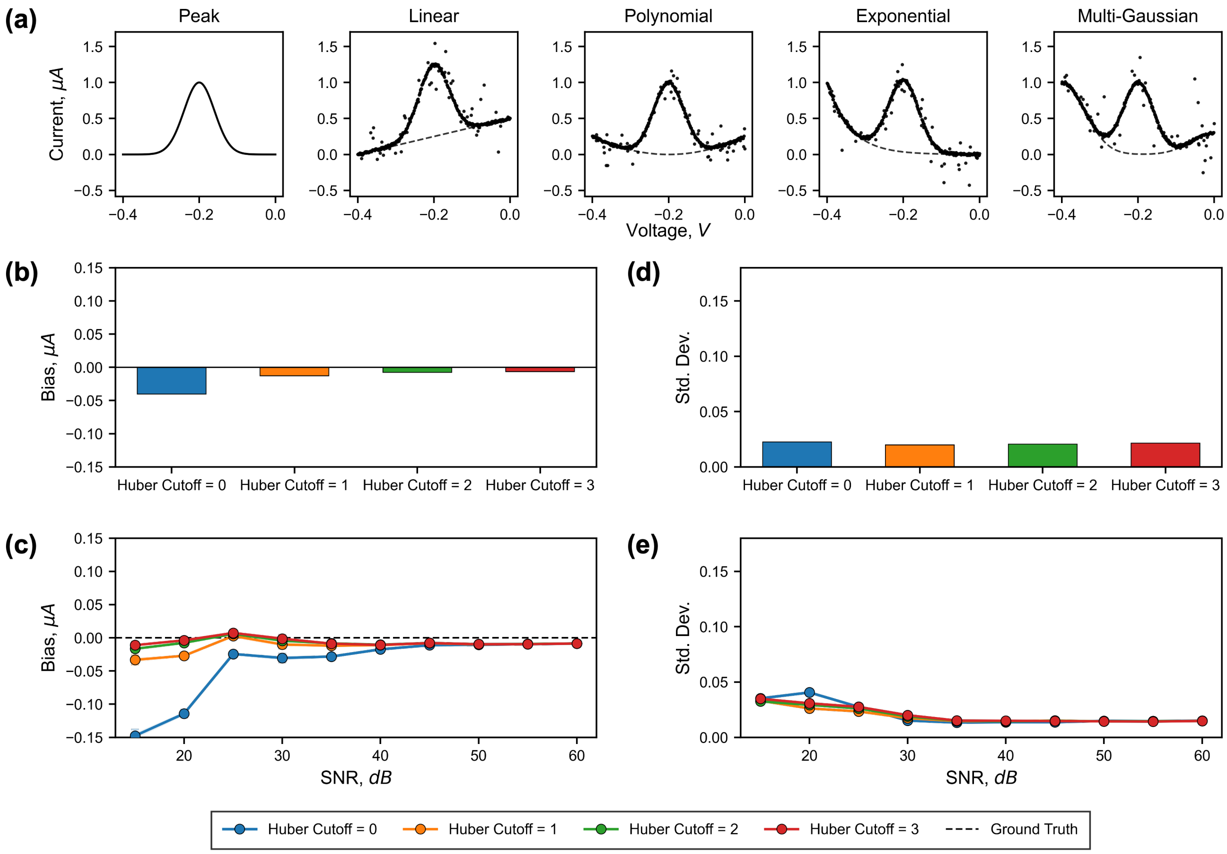


**Figure S10:** Optimization of the Huber cutoff value used by ASWIFT, defined as the residual threshold at which the loss function transitions from a squared-error penalty to a linear penalty. For this analysis, we fixed the rolling median absolute deviation window used for Huber smoothing at 21 samples and fixed the relative-height prominence threshold for peak-region segmentation at 0.3. These parameters were applied to the same simulated panel described in **SI, Fig. 2**, which introduced randomly injected outliers. We found that a Huber cutoff of 2 produced the largest reduction in bias while maintaining comparable standard deviation. Increasing the cutoff to 3 produced little additional improvement, so we selected 2 as the optimal cutoff.

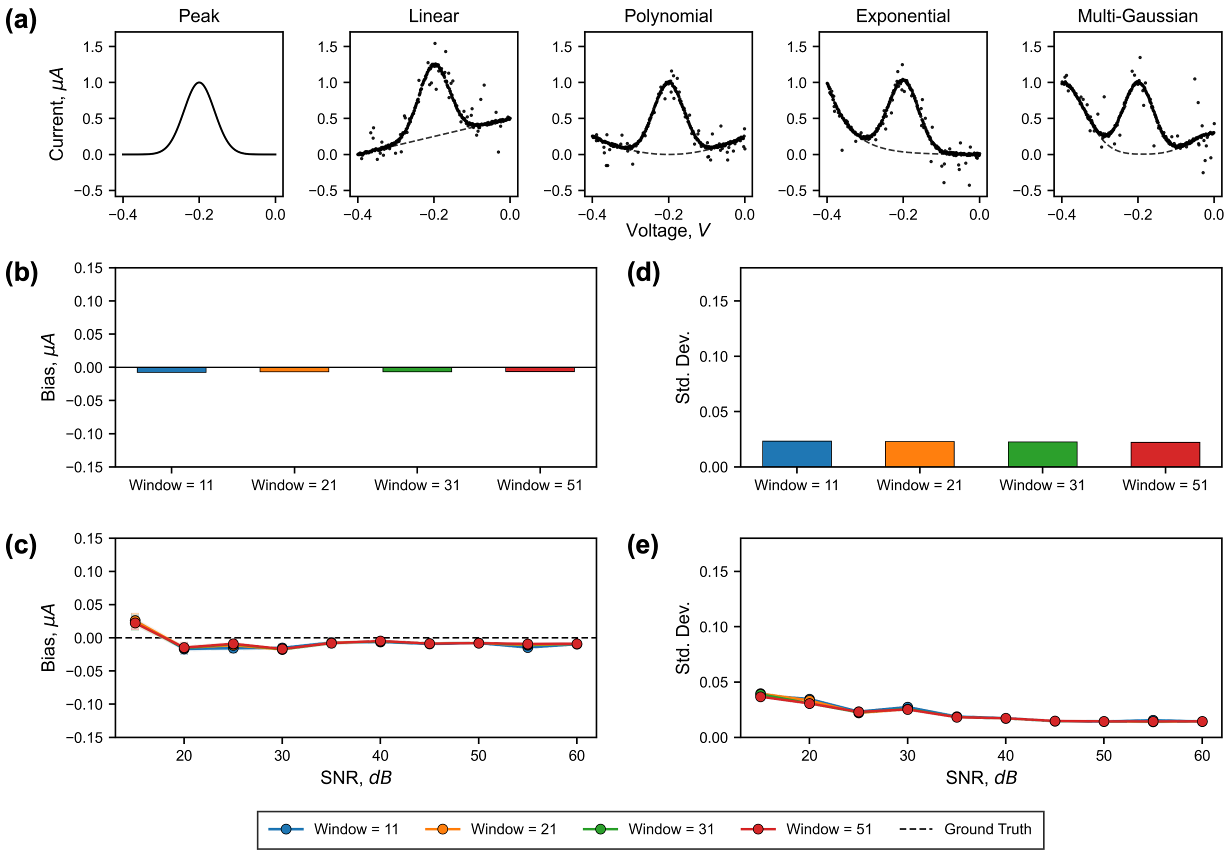


**Figure S11:** Optimization of the local window size used by ASWIFT for Huber down-weighting. For this analysis, outliers were randomly injected into the simulated panel as described in **SI, Fig. 2**, while the Huber cutoff and relative-height prominence threshold were fixed at 2 and 0.3, respectively. Performance varied negligibly across the tested window sizes, so we selected a 21-sample local window for subsequent analyses.

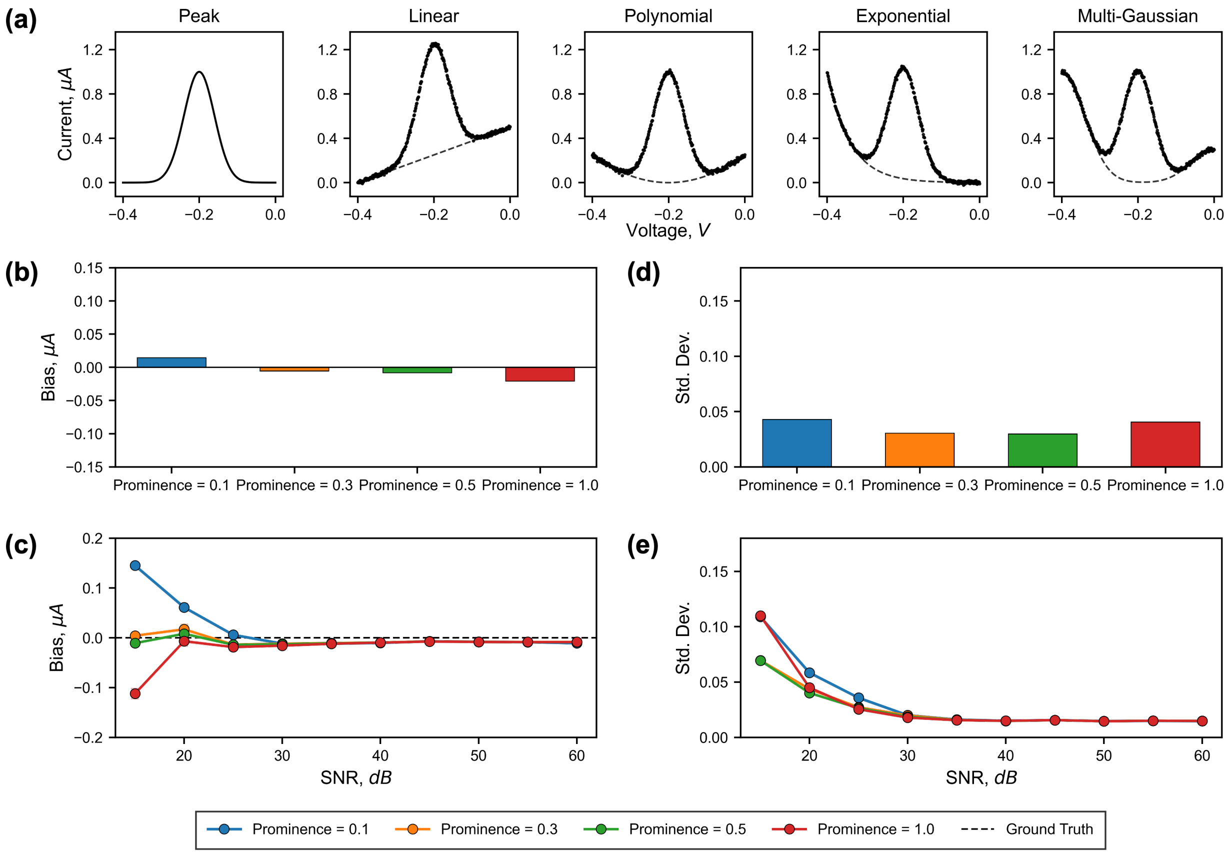


**Figure S12:** Optimization of the relative-height prominence threshold used by ASWIFT for local peak-region segmentation. For this analysis, no additional outliers were injected, and the same simulated panel from **Fig. 2** was used. The Huber cutoff and median absolute deviation window were fixed at 2 and 21 samples, respectively. Low prominence thresholds, such as 0.1, produced positive bias and increased variability, whereas overly high thresholds, such as 1.0, produced negative bias and also increased variability. We therefore selected an intermediate prominence threshold of 0.3, which provided the lowest bias while maintaining higher precision.
