## Supplementary material for "Robust Regularization Enables Automated, Real-Time Square-Wave Voltammetry Signal Quantification": Instructions for installing, configuring, and using ASWIFT.

### ASWIFT User Guide

Version 1.0

#### 1 Setup

ASWIFT can be setup either as a standalone desktop application or as a Python package. The standalone application provides a no-code interface that runs out of the box on modern macOS and Windows systems, while the Python package enables integration into custom analysis pipelines and provides additional customization options.

##### 1.1 Installing the ASWIFT desktop application

Precompiled applications for each supported operating system are available at:

https://github.com/Soh-Lab/aswift/tree/main/release_assets

Choose the appropriate download for your system:

- **macos-arm64**: Apple Silicon (M1, M2, M3, M4, ...)
- **macos-x64**: Intel-based Macs
- **windows-x64**: 64-bit Windows

After downloading, unzip the archive and double-click the **ASWIFT Viewer** application to launch the user interface. Because the application is distributed outside of the Apple App and Microsoft Store, your operating system may display a security warning the first time it is opened. If so, follow the prompts to allow the application to run (e.g., approve the application in **Privacy & Security** on macOS).

##### 1.2 Installing the ASWIFT Python package

The following instructions apply only to users who wish to install ASWIFT as a Python package. Python is not required when using the standalone desktop application. ASWIFT requires Python 3.12 or newer. Install the core package using:

```
$ pip install aswift
```

To install the package together with the interface, pandas CSV helpers, and PalmSens `.pssession` support:

```
$ pip install "aswift[viewer]"
```

PalmSens `.pssession` support depends on `pypalmsens`, which requires the Microsoft .NET 9 Runtime. If ASWIFT reports that it was unable to create a .NET runtime (CoreCLR), install the Microsoft .NET 9 Runtime (https://dotnet.microsoft.com/en-us/download/dotnet/9.0) and restart the application. Example notebooks demonstrating SWV fitting and parameter customization are available at:

https://github.com/Soh-Lab/aswift/tree/main/examples

After installation, activate the same Python environment (e.g., your virtual environment or conda environment) in which ASWIFT was installed, then launch the interface by running:

```
$ aswift-viewer
```

### 2 Usage

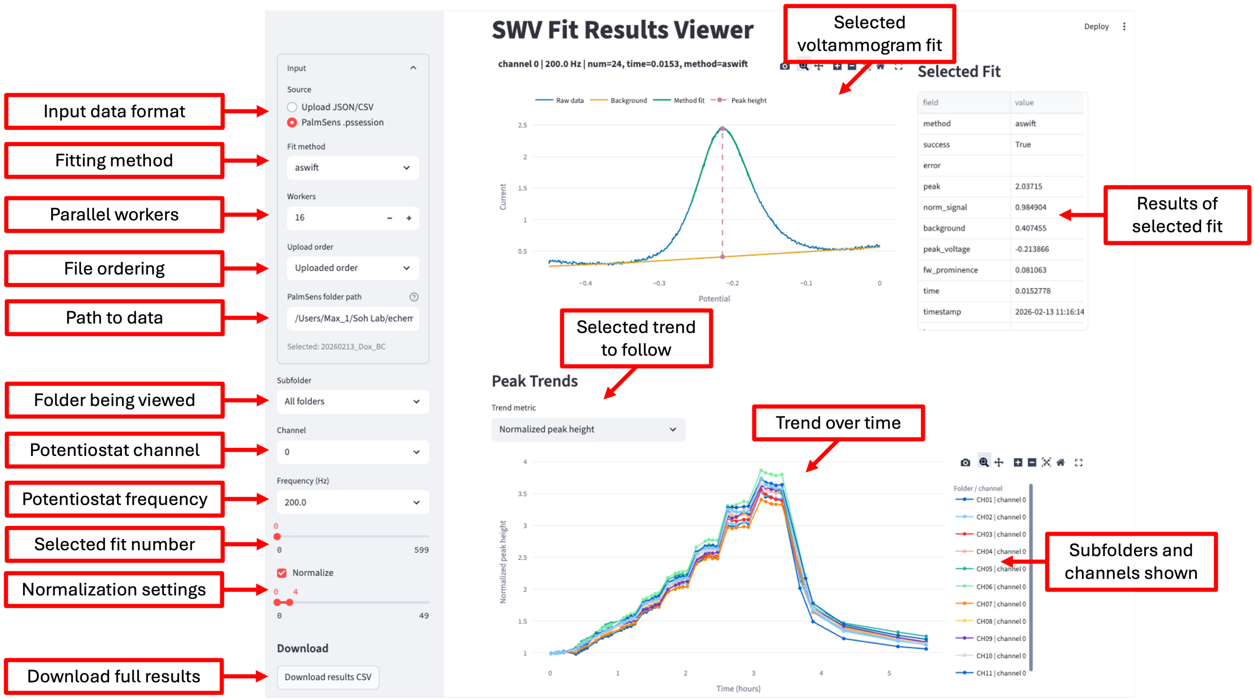


Figure 1: Overview of the ASWIFT Viewer interface. After launching ASWIFT using either the standalone desktop application or the Python package, the user is presented with a similar interface.

#### 2.1 Uploading files

After launching the interface, you will be prompted to either upload one or more SWV data files or specify a directory containing an SWV dataset. ASWIFT currently supports three input file formats. Example files for each format are available at:

https://github.com/Soh-Lab/aswift/tree/main/release_assets

The supported file formats are:

- **.pssession**: Files generated by PalmSens potentiostats. After selecting the `.pssession` input format and specifying a folder containing `.pssession` files (or subfolders containing them), ASWIFT automatically extracts metadata such as frequency, sampling time, and channel information before processing the data. The processed results are saved as a structured `.json` file in the selected directory. If new `.pssession` files are added while the interface is running, ASWIFT automatically detects and processes them in real time.
- **.csv**: ASWIFT supports both simple CSV files containing voltage values in the first column and one or more current traces in subsequent columns, as well as expanded CSV files containing one complete voltammogram per row together with metadata such as channel and frequency for larger datasets. Correctly formatted examples are available in the GitHub release linked above.
- **.json**: Structured output files previously generated by ASWIFT from `.pssession` data. Reloading these files allows previously processed datasets to be visualized and analyzed without repeating the fitting step.

#### 2.2 Processing results

After ASWIFT finishes processing a dataset, the fit and extracted peak height can be inspected for individual voltammograms. Processing can be accelerated on multi-core computers by increasing the number of worker processes. The interface also displays the corresponding time (or measurement) series for peak height, peak location, and full-prominence peak width. The following controls determine what is displayed:

- **Subfolder**: If the input directory contains multiple subfolders of `.pssession` files, select the subfolder of voltammograms and signal trends you wish to visualize.
- **Channel**: Select the channel for which individual voltammogram fits are displayed. The time-series plots initially show all channels, which can be individually hidden or displayed using the plot legend.
- **Frequency**: Select the frequency to visualize. Individual voltammogram fits and time-series plots are displayed for one frequency at a time.
- **Selected fit number**: Select the individual voltammogram to inspect for the chosen subfolder, channel, and frequency. Fit numbers are assigned sequentially based on acquisition time (or upload order when timestamps are unavailable).
- **Normalization settings**: When enabled, peak heights are normalized by the average peak height over the user-specified reference range below. Normalization is performed independently for each channel, frequency, and subfolder using only the selected reference measurements.

After processing is complete, the normalized and raw results can be downloaded as a CSV file containing the extracted signal trends and associated metadata for further analysis.

#### 2.3 Bug reports and feature requests

If you encounter a bug or other problem, please open an issue on the ASWIFT GitHub repository. Questions, feedback, and feature requests are also welcome.
